# Methods for Molecular Recognition Computing

**DOI:** 10.64898/2026.04.03.716328

**Authors:** Sai T. Reddy

## Abstract

The softmax attention mechanism in transformer architectures (Vaswani et al., 2017) is mathematically identical to the Boltzmann distribution governing molecular binding at thermal equilibrium (Boltzmann, 1877). Luce’s Choice Axiom (1959) establishes this function - which we term the convergence equation - as the unique function satisfying five axioms of competitive selection: positivity, normalization, unrestricted domain, rank preservation, and independence of irrelevant alternatives. We show that five additional architecture conditions - discrete intermolecular contacts, bilinear energy decomposition, finite competitor pools, thermal equilibrium, and stochastic selection - are satisfied by at least ten biological molecular recognition systems and together prescribe a complete neural architecture: dual encoders, cross-attention, InfoNCE contrastive training, symmetric loss, learned temperature, and cross-attentive decoder. We term this architecture a Specificity Foundation Model (SFM) and specify it for antibody-antigen, TCR-peptide-MHC, transcription factor-DNA, microRNA-mRNA, enzyme-substrate, CRISPR guide RNA-DNA, drug-target, peptide-MHC, receptor-ligand, and RNA-binding protein-RNA recognition. The first implementation (CALM; Lee et al., 2026) achieves antibody-antigen retrieval from approximately 4,000 training pairs with ∼100,000-fold greater data efficiency than comparable contrastive architectures trained without the physics derivation. We classify this as Level 3 architecture-physics alignment and derive three further theoretical results: an exponential scaling law for retrieval accuracy as a function of training data diversity (the MRC scaling law), a two-parameter affinity calibration framework connecting contrastive scores to binding free energies, and a hybrid recursive learning framework for cross-modal reinforcement learning with orthogonal verification. The failure conditions of the framework are analyzed in terms of the validity of equilibrium thermodynamics for molecular binding and the convergence properties of gradient-based parameter estimation.

## §1. Introduction

Computational approaches to molecular biology have progressed through several paradigms over sixty years. Molecular mechanics force fields established the mathematical vocabulary of energy minimization (Lifson & Warshel, *J. Chem. Phys*., 1968; Cornell *et al., J. Am. Chem. Soc*., 1995). The CASP competitions created a rigorous benchmarking culture for structure prediction (Moult *et al., Proteins*, 1995). The Rosetta framework demonstrated computational design of novel functional proteins (Kuhlman *et al., Science*, 2003). AlphaFold2 solved single-chain protein structure prediction (Jumper *et al., Nature*, 2021). Protein language models learned structural features from evolutionary sequence data (Rives *et al., PNAS*, 2021; Lin *et al., Science*, 2023). AlphaFold3 extended structure prediction to heterogeneous molecular complexes (Abramson *et al., Nature*, 2024). Deep learning design tools — RFdiffusion (Watson *et al., Nature*, 2023), ProteinMPNN (Dauparas *et al., Science*, 2022), BoltzGen (Stark *et al., bioRxiv*, 2025) — enabled de novo protein binder generation. Two Nobel Prizes in Chemistry (2013, 2024) trace their lineage to this progress.

Throughout this development, one problem has remained unsolved: molecular recognition. Predicting which molecules bind which other molecules — which antibody recognizes which antigen, which transcription factor binds which DNA site, which drug binds which protein target — has not been achieved by structure-first methods. The dominant paradigm assumes that the path to specificity passes through structure: predict molecular geometry, then infer binding from predicted structures. This logic is productive for folding and design but introduces mathematical difficulties for recognition, including ill-posed inverse mappings, proxy loss functions, and information-theoretic expansion through high-dimensional intermediate representations (§9).

A separate line of work established that molecular sequence contains sufficient information for specificity prediction, at least within defined molecular neighborhoods. Deep mutational learning demonstrated that neural networks trained on display screening data could predict antibody-antigen specificity from sequence (Mason *et al., Nat. Biomed. Eng*., 2021; Taft *et al., Cell*, 2022). Cross-modal contrastive learning — particularly CLIP (Radford *et al., ICML*, 2021) and its extensions — provided the architectural framework for learning shared embedding spaces between two molecular modalities. The noise-contrastive estimation framework (Gutmann & Hyvärinen, *AISTATS*, 2010; Oord *et al., arXiv*, 2018) provided the training objective. The connection between softmax attention and energy-based models was noted by Ramsauer *et al*. (*ICLR*, 2021) in the context of Hopfield networks. What was missing was the theoretical framework connecting these components to the physics of molecular binding.

This paper provides that framework. The central result is that the Boltzmann distribution governing molecular binding at thermal equilibrium (Boltzmann, 1877) and the softmax attention mechanism in transformers (Vaswani *et al*., 2017) are the same function, and that Luce’s Choice Axiom (Luce, 1959) proves this function is unique. Combined with five architecture conditions satisfied by molecular recognition systems, this identity prescribes a complete neural architecture — a Specificity Foundation Model (SFM) — across at least ten biological domains. The theory extends to scaling predictions, affinity calibration, and cross-modal reinforcement learning.

We build on two published works: computational convergence of adaptive immunity and artificial intelligence (Reddy, *bioRxiv*, 2026), which proved the Boltzmann-softmax identity for adaptive immune recognition, and Cross-attention Adaptive immune receptor-Antigen Language Model, CALM (Lee *et al., bioRxiv*, 2026), which demonstrated the prescribed architecture for antibody-antigen sequence-to-specificity prediction.

The present paper consolidates the complete mathematical framework and presents the methods for molecular recognition computing (MRC). Section 2 derives the convergence equation and its uniqueness. Section 3 derives the SFM architecture from five physical conditions. Section 4 classifies architecture-physics alignment into three levels. Section 5 specifies ten domain-specific SFMs. Sections 6–8 present three further theoretical results: the MRC scaling law (§6), a two-parameter affinity calibration framework connecting contrastive scores to binding free energies (§7), and hybrid recursive learning with orthogonal verification (§8). Section 9 compares the sequence-first and structure-first paradigms. Section 10 analyzes the conditions under which the framework can fail. Section 11 concludes. Full proofs are provided in Supplementary Notes S1–S5.

## §2. The Convergence Equation

Molecular recognition reduces to a single mathematical problem: convert compatibility scores into selection probabilities. An antibody paratope has compatibility scores with every epitope on every antigen surface. A transcription factor has compatibility scores with every candidate DNA binding site in the genome. A transformer’s query token has dot-product scores with every key token in the sequence. All three systems face the same problem. We require five properties of any function that solves it. Each property eliminates candidate functions. After five eliminations, one function remains.

### 2.1 The Five Axioms of Competitive Selection

We state five properties that any physically meaningful selection function must satisfy. These are not modeling choices. They are constraints imposed by the physics of competitive molecular binding at thermal equilibrium.

1. **Positivity**. Every candidate has non-negative selection probability: *P*(*j*) > 0 for all *j* and all score vectors. At non-zero temperature, every molecular interaction has non-zero probability. A selection function that assigns exactly zero probability to a reachable state contradicts thermodynamics.
2. **Normalization**. Probabilities sum to one: ∑_*j*_ *P* (*j*) = 1. The agent binds exactly one target at a time, or equivalently, the probability distribution over binding states sums to unity.

Combined with positivity, normalization forces the selection function to have the form

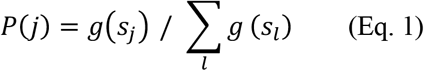

where *g*: ℝ → ℝ^*+*^ is a strictly positive function. The question reduces to: what is *g*?

1. **Unrestricted domain**. The function *g* must accept any real-valued input: dom(*g*) = ℝ. In molecular binding, energies span the full real line — favorable interactions have negative *ΔG*, unfavorable interactions have positive *ΔG*. In transformer attention, dot-product scores are unrestricted. Any function that requires non-negative inputs is disqualified.
2. **Rank preservation**. Higher scores yield higher probabilities: if *s*_*A*_ > *s*_*B*_ then *P*(*A*) > *P*(*B*). A selection function that does not respect the ordering of compatibility scores is not computing selection — it is introducing noise. This requires *g* to be strictly increasing.
3. **Independence of irrelevant alternatives (IIA)**. The ratio of selection probabilities between any two candidates depends only on their own scores, not on what other candidates are present in the pool:

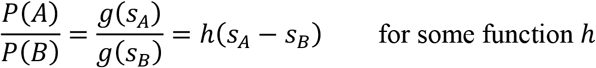

This is the decisive property. It states that the relative preference between antibody binding target *A* versus target *B* does not change when a third target *C* enters or leaves the pool. In economics, this is Luce’s Choice Axiom (Luce, *Individual Choice Behavior*, 1959). In physics, it follows from the fact that the binding free energy between two molecules is a property of that pair, independent of other molecules in solution.

### 2.2 The Unique Survivor

Properties 1–4 constrain *g* to be a strictly positive, strictly increasing function defined on all of ℝ. Many functions satisfy these four properties: the exponential *e*^*s*^, the sigmoid 1/(1 + *e*^−*s*^), the function arctan(*s*) + *π*/2, among others. Property 5 eliminates all but one.

Consider the sigmoid: *σ*(*s*) = (1/ + exp(−*s*)). It satisfies Properties 1–4. But the ratio *σ*(*s*_*A*_)/*σ*(*s*_*B*_) expands to

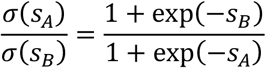

This ratio depends on the absolute magnitudes of *s*_*A*_ and *s*_*B*_, not only on the difference *s*_*A*_ − *s*_*B*_. Sigmoid fails IIA.

The exponential function satisfies IIA exactly:

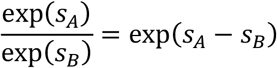

The denominator cancels. The ratio depends only on the score difference. No other function family achieves this while simultaneously satisfying Properties 1–4. The full elimination — including polynomial, periodic, power-law, and composite candidates — is given in Supplementary Note S1. Each family fails at least one property.

Luce proved in 1959 that the exponential normalization function is the unique function satisfying Properties 1–5 simultaneously (Luce, *Individual Choice Behavior*, 1959). We refer to this result as the **Luce uniqueness theorem**. The unique survivor is:

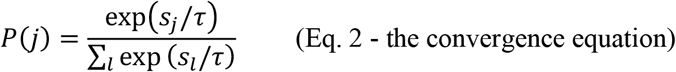

This is the softmax function. The parameter *τ* (temperature) controls the sharpness of selection: as *τ* → 0, selection concentrates on the highest-scoring candidate; as *τ* → ∞, selection becomes uniform. The functional form is determined by the five axioms. Temperature is the only free parameter.

### 2.3 Five Independent Derivations

Equation 2 has been derived at least five times independently across 150 years.

1. **Statistical mechanics**. Boltzmann derived the distribution from maximum entropy at thermal equilibrium: the most probable macrostate of a system in thermal contact with a heat bath assigns occupation probabilities proportional to exp(−*E*/*k*_*B*_*T*) (Boltzmann, *Wien. Ber*., 1877).
2. **Information theory**. Shannon’s entropy *H* = −∑*p*_*i*_ log *p*_*i*_ is Boltzmann’s entropy with *k*_*B*_ = 1. The distribution that maximizes Shannon entropy subject to a mean constraint is the Boltzmann distribution — Equation 2. Shannon recognized the mathematical identity and adopted the term “entropy” from thermodynamics (Shannon, *Bell System Technical Journal*, 1948).
3. **Maximum entropy inference**. Jaynes unified the Boltzmann and Shannon derivations by proving that the exponential distribution is the uniquely rational probability assignment under incomplete information: given any set of constraints, the distribution that is maximally noncommittal with respect to missing information is Equation 2. He predicted explicitly that this form would appear in any domain involving selection under uncertainty (Jaynes, *Phys. Rev*., 1957).
4. **Mathematical psychology**. Luce proved that the exponential normalization is the unique function satisfying the axioms of rational choice — in particular, the independence of irrelevant alternatives. No other function satisfies this axiom while simultaneously satisfying positivity, normalization, unrestricted domain, and rank preservation (Luce, *Individual Choice Behavior*, 1959). McFadden applied this result to consumer choice in econometrics (McFadden, *Frontiers in Econometrics*, 1974; Nobel Prize in Economics, 2000).
5. **Machine learning**. Vaswani *et al*. implemented Equation 2 as scaled dot-product attention — softmax 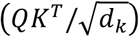 — in the transformer architecture (Vaswani *et al., NeurIPS*, 2017). The temperature parameter 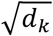 plays the same role as *k*_*B*_*T* in the Boltzmann distribution

#### Molecular biology

The present work and its companion (Reddy, *bioRxiv*, 2026) is not an independent derivation, it has been established that the Boltzmann distribution (convergence equation) governs molecular recognition across all biological systems at thermal equilibrium. Rather the present work establishes that combined with the five architecture conditions (Section 3), this identity prescribes a complete neural architecture (Specificity Foundation Model) for molecular specificity prediction. The contribution is not the equation — it is the proof that the equation prescribes the architecture.

Each community encountered the same problem — converting scores into selection probabilities — and arrived at the same solution. What Boltzmann called a partition function, Shannon called a maximum entropy distribution, Luce called a choice denominator, and Vaswani called an attention normalizer are the same mathematical object.

### 2.4 Application to Molecular Binding

At thermal equilibrium, the probability that an agent molecule *A* selects target *j* from a pool of competitors follows the Boltzmann distribution:

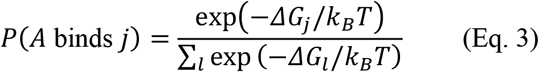

where *ΔG*_*j*_ is the binding free energy between *A* and target *j, k*_*B*_ is Boltzmann’s constant, and *T* is absolute temperature.

Defining a dimensionless compatibility score *s*_*j*_ = −*ΔG*_*j*_/*k*_*B*_*T*, Equation 3 becomes *P*(*j*) = softmax(*s*_*j*_), which is Equation 2 with *τ* = 1 in natural units. What molecular biology calls a binding probability and what machine learning calls an attention weight are the same mathematical object. The identity is exact — algebraic, not approximate.

Every *K*_*d*_ measurement ever made by surface plasmon resonance, isothermal titration calorimetry, or any other binding assay is a measurement of the Boltzmann probability ratio between bound and unbound states. The convergence equation has been verified experimentally across every domain of chemistry and biophysics for over 150 years.

### 2.5 Prior Art

The identity between the Boltzmann distribution and softmax has been recognized since Gibbs formalized statistical mechanics (Gibbs, 1902). Bridle introduced the term “softmax” in neural networks (Bridle, *NIPS*, 1989). Sutton and Barto use “Boltzmann exploration” as standard terminology in reinforcement learning (Sutton & Barto, 1998). The Artificial Immune Systems field has explored immune-computation connections since the 1990s, through algorithmic rather than mathematical frameworks (Forrest *et al*., 1994; de Castro & Timmis, 2002). Franke and Degen provide a tutorial deriving softmax from multiple perspectives including maximum entropy (Franke & Degen, *arXiv*, 2023). Ramsauer *et al*. proved that transformer attention is equivalent to the update rule of a modern Hopfield network, establishing a connection between energy-based models and attention (Ramsauer *et al., ICLR*, 2021). Anandkumar *et al*. developed physics-informed neural operators that incorporate physical constraints into architectures for PDE systems (Azizzadenesheli *et al., Nat. Rev. Phys*., 2024).

Our contribution is not the Boltzmann-softmax identity. It is the proof that the identity, combined with the five architecture conditions (Section 3), prescribes a complete neural architecture for molecular specificity prediction across ten biological domains — and that this prescription extends to the training objective, loss symmetry, temperature, decoder, affinity calibration, and scaling behavior. We call Equation 2 **the convergence equation** because it is the point where six fields converge on the same function.

## §3. Derivation of the Specificity Foundation Model Architecture

The convergence equation (Eq. 2) governs any system at thermal equilibrium — this is unconditional. But the full identity between the convergence equation and transformer attention, which yields a Specificity Foundation Model (SFM), requires five additional conditions. A molecular recognition system satisfying all five admits a physics-derived neural architecture in which every design choice follows from the governing equation.

### 3.1 The Five Architecture Conditions

1. **Discrete intermolecular contacts**. Binding occurs through a finite set of contacts between specific positions on two molecules. Antibody CDR residues contact antigen epitope residues. Transcription factor amino acids contact DNA nucleotides. Guide RNA nucleotides contact protospacer nucleotides. This discreteness enables positional encoding: each contact position contributes independently to the binding energy.
2. **Bilinear energy decomposition**. The binding energy decomposes as a sum of pairwise contributions between agent and target positions:

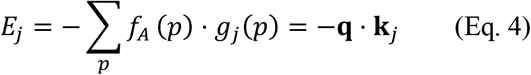

where **q** encodes agent features and **k**_*j*_ encodes target features. Substituting into the convergence equation:

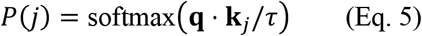

This is the transformer attention equation. The bilinear decomposition is the bridge between thermodynamics and the query-key dot product. In nucleic acid systems (TF–DNA, miRNA–mRNA, CRISPR), bilinearity is **given**: the energy decomposes exactly by position in raw sequence space, as encoded by position weight matrices (Stormo, *Bioinformatics*, 2000) or nearest-neighbor thermodynamic models (SantaLucia, *PNAS*, 1998). In protein systems (antibody–antigen, TCR–pMHC, enzyme–substrate), bilinearity is **emergent**: the energy depends on 3D contacts that are not directly readable from the linear sequence, so the bilinear decomposition must be learned from data by the encoder. This distinction — given versus emergent bilinearity — predicts measurably different scaling behavior across domains (Section 6). Full verification of all five conditions across all ten domains is provided in Supplementary Note S2.
3. **Finite competitor pools**. The agent selects among a finite number of candidate targets, yielding a finite softmax denominator. This is satisfied by all biological recognition systems operating within a cell, tissue, or organism.
4. **Thermal equilibrium**. The system operates at a well-defined temperature. All ten molecular recognition systems operate in aqueous solution at approximately 310 K, where *k*_*B*_*T* ≈ 0.6 kcal/mol. The equilibrium assumption is standard for binding affinity measurements and is explicitly validated by SPR and ITC kinetics.
5. **Stochastic selection**. Binding is probabilistic at physiological temperature, requiring softmax rather than argmax. At *T* = 310 K, the thermal energy *k*_*B*_*T* ≈ 0.6 kcal/mol is comparable to the energy differences between binding partners, ensuring that selection is stochastic rather than deterministic.

### 3.2 The Prescribed Architecture

The convergence equation and the five architecture conditions together prescribe the following components:

#### Dual encoders

Two separate encoders map agent and target molecules into a shared embedding space. The agent encoder produces query vectors **q**; the target encoder produces key vectors **k**. This follows from Condition 2: the bilinear energy decomposition requires independent feature representations for each molecular partner. The dual-encoder architecture is established in cross-modal representation learning (Radford *et al., ICML*, 2021). In practice, each encoder is initialized from a pre-trained language model appropriate to the molecular modality: protein language models such as ESM-2 (Lin *et al., Science*, 2023) or domain-specific models such as AntiBERTy for antibody sequences (Ruffolo *et al., Patterns*, 2022), nucleic acid language models such as DNABERT-2 (Zhou *et al., iScience*, 2024) or RNA-FM for RNA sequences, and molecular language models for small-molecule drugs. The pre-training paradigm follows the masked language modeling approach introduced by BERT (Devlin *et al., NAACL*, 2019) and extended to protein sequences by ESM (Rives *et al., PNAS*, 2021). CALM uses AntiBERTy for the antibody encoder and ESM-2 for the antigen encoder (Lee *et al., bioRxiv*, 2026).

#### Cross-attention

The compatibility score between agent *i* and target *j* is computed as the dot product **q**_*i*_ ⋅ **k**_*j*_, and the selection probability follows from Equation 5. This is scaled dot-product attention (Vaswani *et al., NeurIPS*, 2017). It is not an architectural choice — it is prescribed by the bilinear energy decomposition and the convergence equation.

#### Multi-head attention

In protein systems, binding involves multiple partially independent contact regions — for example, the three CDR loops of an antibody, each contacting different parts of the antigen surface (Reddy, *bioRxiv*, 2026). Multi-head attention, where *H* parallel attention heads compute independent compatibility scores that are concatenated (Vaswani *et al., NeurIPS*, 2017), corresponds to this partial independence. In nucleic acid systems, position-specific contacts are closer to fully independent, and multi-head attention captures positional contributions. The correspondence between heads and contact regions is structural rather than exact (Reddy, *bioRxiv*, 2026).

### 3.3 The Training Objective

The training objective is derived from the convergence equation by maximum likelihood estimation. A molecular agent encounters a pool containing its correct target *f* and *M* non-cognate competitors {*c*_1_, …, *c*_*M*_}. The selection probability is:

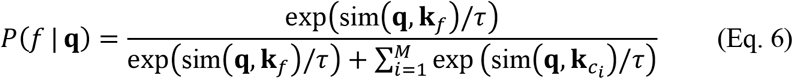

Maximizing this probability across all training pairs and taking the negative logarithm yields:

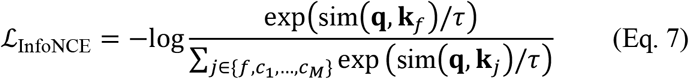

This is the InfoNCE contrastive loss (Oord *et al., arXiv*, 2018), which extends noise-contrastive estimation (Gutmann & Hyvärinen, *AISTATS*, 2010) to the multi-class setting. It was not selected from a menu of objectives — it is the negative log-likelihood of the convergence equation applied to the training scenario. In the immune system, this corresponds to the clonal selection probability: a lymphocyte correctly selecting a foreign antigen over self-antigens (Reddy, *bioRxiv*, 2026). The full derivation, including elimination of alternative objectives, is in Supplementary Note S3.

#### Why not other contrastive losses

Triplet loss (Schroff *et al., CVPR*, 2015) enforces a margin between positive and negative similarities but does not model the full competitor pool — it violates the normalization axiom by considering only one negative at a time. Binary cross-entropy treats each pair independently, ignoring the competitive context — it violates IIA by not conditioning on the full alternative set. Margin-based losses introduce a hyperparameter (the margin) that has no physical interpretation in the convergence equation. Each alternative violates at least one of the five axioms. InfoNCE is the only contrastive objective consistent with the convergence equation.

### 3.4 Symmetric Loss

Binding free energy is a property of the complex, not of either partner individually: *ΔG*_*AB*_ = *ΔG*_*BA*_. This physical symmetry requires the training loss to be bidirectional. The symmetric loss averages the InfoNCE loss computed in both directions:

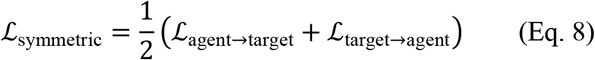

where ℒ_agent→target_ uses agent embeddings as queries and target embeddings as keys, and ℒ_target→agent_ reverses the roles. This follows the formulation of CLIP (Radford *et al., ICML*, 2021) and is used in CALM (Lee *et al., bioRxiv*, 2026). It is derived here from binding symmetry rather than adopted as a heuristic. Within each mini-batch, all non-cognate pairs serve as the competitor pool (in-batch negatives).

### 3.5 Hard Negatives

Not all competitors are equally informative. A non-binding molecule that shares no features with the true target provides a trivially discriminable negative. A molecule that shares partial features — surface complementarity without the correct charge distribution, seed-region match without 3’ supplementary pairing — provides a hard negative that forces the model to learn fine-grained discrimination.

In the immune system, thymic negative selection mediated by the autoimmune regulator AIRE (Anderson *et al., Science*, 2002) presents self-antigens as hard negatives during T cell maturation; the correspondence between AIRE-mediated selection and hard negative mining was established in Reddy (*bioRxiv*, 2026). This principle generalizes across domains: in tSFM, paralogous transcription factors with similar binding motifs serve as hard negatives. In crisprSFM, off-target sites with seed-region matches are hard negatives (Hsu *et al., Nat. Biotechnol*., 2013). In TCR-SFM, cross-reactive peptides that bind multiple TCRs serve as hard negatives. In each case, the hard negatives are defined by the domain-specific biology, but their computational role is identical: they populate the denominator of Equation 6 with informative competitors that sharpen the embedding geometry. Hard negative mining strategies for contrastive learning are reviewed in Robinson *et al*. (*NeurIPS*, 2021).

### 3.6 Temperature

The parameter *τ* in the convergence equation has a physical interpretation: it corresponds to *k*_*B*_*T*, the thermal energy. In standard transformer attention, 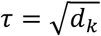 is set by the key dimension (Vaswani *et al., NeurIPS*, 2017). In SFMs, *τ* is a learned parameter, following CLIP (Radford *et al., ICML*, 2021) and CALM (Lee *et al., bioRxiv*, 2026). The prediction is that *τ*_learned_ should recover a value proportional to the physical temperature of the system, up to a unit conversion determined by the embedding geometry. This prediction is directly testable through the affinity calibration framework (Section 7). We call this **temperature recovery**.

At low *τ*, the convergence equation concentrates probability on the highest-scoring target (sharp selection). At high *τ*, probability distributes more uniformly across competitors (permissive selection). Biologically, this corresponds to the stringency of molecular selection: a tightly regulated system (e.g., CRISPR on-target recognition) operates at effectively lower temperature than a system tolerating broader cross-reactivity (e.g., innate immune receptors).

### 3.7 The Cross-Attentive Decoder

The components above — dual encoders, cross-attention, InfoNCE, symmetric loss, learned temperature — constitute the discriminative SFM: given an agent, retrieve the most likely target (and vice versa). The decoder extends this to generation: given a target, generate a novel agent sequence (or vice versa).

The convergence equation prescribes one invariant: the decoder conditions on encoder states through cross-attention, where decoder-side representations serve as queries and encoder-side representations serve as keys and values:

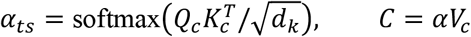

The cross-attention weight *α*_*ts*_ is the probability that position *t* of the generated sequence corresponds to position *s* of the source — this is the convergence equation applied at the residue-contact level. This cross-attention interface is shared by all decoder variants.

Two decoder architectures are compatible with this interface. The **autoregressive (AR) decoder** generates tokens sequentially, conditioning each position on all previously generated positions (Raffel *et al., JMLR*, 2020). This is the conservative choice: well-understood, with many years of engineering from neural machine translation and language modeling. The **discrete diffusion (DD) decoder** generates all positions in parallel through iterative denoising, starting from a fully masked sequence and progressively unmasking positions in order of model confidence (Austin *et al., NeurIPS*, 2021; Sahoo *et al., arXiv*, 2024). The DD decoder is the more physics-aligned choice: in the Boltzmann framework, binding occupancy at each target position is resolved simultaneously at equilibrium, not sequentially. The autoregressive factorization *P*(*x*_1_)*P*(*x*_2_|*x*_1_) ⋯ *P*(*x*_*L*_|*x*_<*L*_) imposes a left-to-right ordering that the binding physics does not have. For nucleic acid domains (tSFM, miR-SFM, crisprSFM), where positional independence approximately holds, the DD decoder is expected to be particularly well-matched. The choice between AR and DD is empirical and domain-dependent; the cross-attention interface to the encoders is invariant.

Bidirectional generation is enabled by task tokens: a ⟨to_target⟩ token conditions the decoder to generate a target sequence given an agent, and a ⟨to_agent⟩ token reverses the direction. This mirrors multilingual neural machine translation, where a single decoder with a target-language token enables many-to-many translation (Johnson *et al., TACL*, 2017).

The decoder is trained in three stages, following the approach demonstrated in CALM (Lee *et al., bioRxiv*, 2026): (1) contrastive pre-training of the dual encoders (InfoNCE), (2) decoder warm-up with frozen encoders, and (3) joint fine-tuning with a combined loss ℒ = *λ*ℒ_InfoNCE_ + (1 − *λ*)ℒ_decoder_, where *λ* is annealed from 1.0 to 0.2 to preserve the contrastive embedding geometry while enabling conditional generation.

With a trained decoder, an SFM is a complete sequence-to-sequence generative system: it retrieves binding partners from sequence and generates novel binding sequences from sequence, without requiring structural input at inference. Structural models such as AlphaFold3 (Abramson *et al., Nature*, 2024) and Boltz-2 (Wohlwend *et al., arXiv*, 2025) serve as training data infrastructure — pseudo-labeling oracles that generate synthetic binding pairs for contrastive training.

### 3.8 Summary: What Is Prescribed and What Is Not

The convergence equation and the five architecture conditions prescribe the following:

| Prescribed by theory | Domain-specific (learned from data) |
| --- | --- |
| Softmax attention (convergence equation) | Encoder weights |
| Dual-encoder architecture (bilinear decomposition) | Embedding dimensions |
| InfoNCE training objective (maximum likelihood) | Batch size, learning rate |
| Symmetric loss (binding symmetry) | Hard negative mining strategy |
| Learned temperature (physical $k_B T$) | Domain-specific pooling masks |
| Cross-attentive decoder interface (Boltzmann contact probability) | Decoder type (AR vs DD), depth, width |

The left column is determined by the mathematics. The right column is determined by the data and the domain. Building an SFM for a new molecular recognition domain requires specifying the right column — choosing encoders, curating training data, designing domain-appropriate hard negatives — while the left column remains invariant. CALM demonstrates this for antibody–antigen (Lee *et al., bioRxiv*, 2026). The expansion from one SFM to ten is engineering, not exploration: the architecture is fixed; the domain-specific configuration is the work.

## §4. Classification of Architecture-Physics Alignment

The relationship between a neural architecture and the physical system it models can be classified into three levels of increasing mathematical stringency.

### Level 1: Analogy

The architecture is inspired by a physical or biological system, but the mathematical form differs substantially. McCulloch-Pitts neurons (McCulloch & Pitts, *Bulletin of Mathematical Biophysics*, 1943) were inspired by biological neurons; the mathematical operations (thresholded linear sums) do not implement the electrochemical dynamics of neuronal firing. Artificial immune system algorithms (Forrest *et al*., 1994; de Castro & Timmis, 2002) borrow the vocabulary of clonal selection and somatic hypermutation; the algorithms do not implement the Boltzmann distribution governing these processes. At Level 1, the physical system provides motivation but does not constrain the architecture.

### Level 2: Structural correspondence

The architecture shares organizational features with the physical system, and the mathematics is related but not identical. Convolutional neural networks share the hierarchical, spatially local, translation-invariant processing of the mammalian visual cortex (Hubel & Wiesel, *Journal of Physiology*, 1962; LeCun *et al., PNAS*, 1989). Physics-informed neural networks incorporate PDE constraints as loss terms or architectural priors (Raissi *et al., Journal of Computational Physics*, 2019). Physics-informed neural operators learn solution operators for PDE families while incorporating physical constraints (Azizzadenesheli *et al., Nat. Rev. Phys*., 2024). Ramsauer *et al*. proved that transformer attention is mathematically equivalent to the update rule of a modern Hopfield network with continuous states (Ramsauer *et al., ICLR*, 2021), establishing a Level 2 correspondence between energy-based models and attention. At Level 2, the physical system constrains or informs the architecture, but the architecture does not implement the governing equation of the physical system.

### Level 3: Mathematical identity

The architecture implements the governing equation of the physical system it models. Softmax attention is the Boltzmann distribution (§2). InfoNCE is the maximum likelihood estimator of the Boltzmann selection probability (§3.3). The learned temperature corresponds to the physical temperature (§3.6). The symmetric loss follows from binding free energy symmetry (§3.4). The cross-attentive decoder implements Boltzmann-weighted contact probabilities (§3.7). At Level 3, the architecture is not informed by physics — it is derived from physics. The model performs parameter estimation in a known equation rather than function approximation in an unknown function class.

SFMs constitute, to our knowledge, one of the first Level 3 architecture-physics alignment in deep learning. The distinction from Level 2 is precise: physics-informed neural networks and neural operators incorporate physical constraints into architectures that were designed empirically; SFMs implement an architecture that was derived from the governing equation. The constraints in Level 2 are additive (physics loss terms added to a data-driven objective). The identity in Level 3 is constitutive (the architecture is the equation).

### Consequences for learning

The classification predicts qualitatively different learning regimes. At Level 1, the model must discover both the functional form and the parameters from data — standard function approximation. At Level 2, the model is constrained toward the correct functional form but must still learn it — constrained function approximation. At Level 3, the functional form is given by construction; the model must learn only the parameters that populate the known equation — parameter estimation. Parameter estimation in a known equation is a strictly easier problem than function approximation in an unknown function class, with well-characterized convergence properties (Robbins & Monro, *Annals of Mathematical Statistics*, 1951). This predicts that Level 3 architectures should require less training data to achieve equivalent performance, and should exhibit different scaling behavior, compared to Level 1 or Level 2 architectures applied to the same task. These predictions are developed quantitatively in Section 6.

CALM provides preliminary evidence consistent with this classification. Trained on approximately 4,000 antibody-antigen binding pairs from the PDB (Dunbar *et al., Nucleic Acids Res*., 2014), CALM achieves retrieval performance comparable to in-distribution performance of CLIP (Radford *et al., ICML*, 2021), which uses the same class of contrastive dual-encoder architecture but was trained on 400 million image-text pairs (Lee *et al., bioRxiv*, 2026). The approximately 100,000-fold difference in training data is consistent with the prediction that Level 3 alignment reduces the learning problem from function approximation to parameter estimation. These observations are suggestive but do not constitute proof of the classification; systematic scaling experiments across multiple domains (Section 6) are required to test the prediction rigorously.

## §5. Domain-Specific Specificity Foundation Models

The convergence equation (Boltzmann distribution) governs any molecular binding event at thermal equilibrium. The five architecture conditions (§3.1) are satisfied by any system in which two molecular species interact through discrete contacts with bilinear energy decomposition in a finite competitor pool. These conditions are not specific to any particular biological domain. We have verified them in detail for at least ten molecular recognition systems spanning adaptive immunity, gene regulation, RNA biology, enzymology, genome editing, pharmacology, and cell signaling. For each, we specify a domain-specific SFM: the same base architecture (§3) configured with appropriate encoders, pooling masks, and training data.

Table 1 summarizes the ten domains against the five architecture conditions.

**Table 1.**
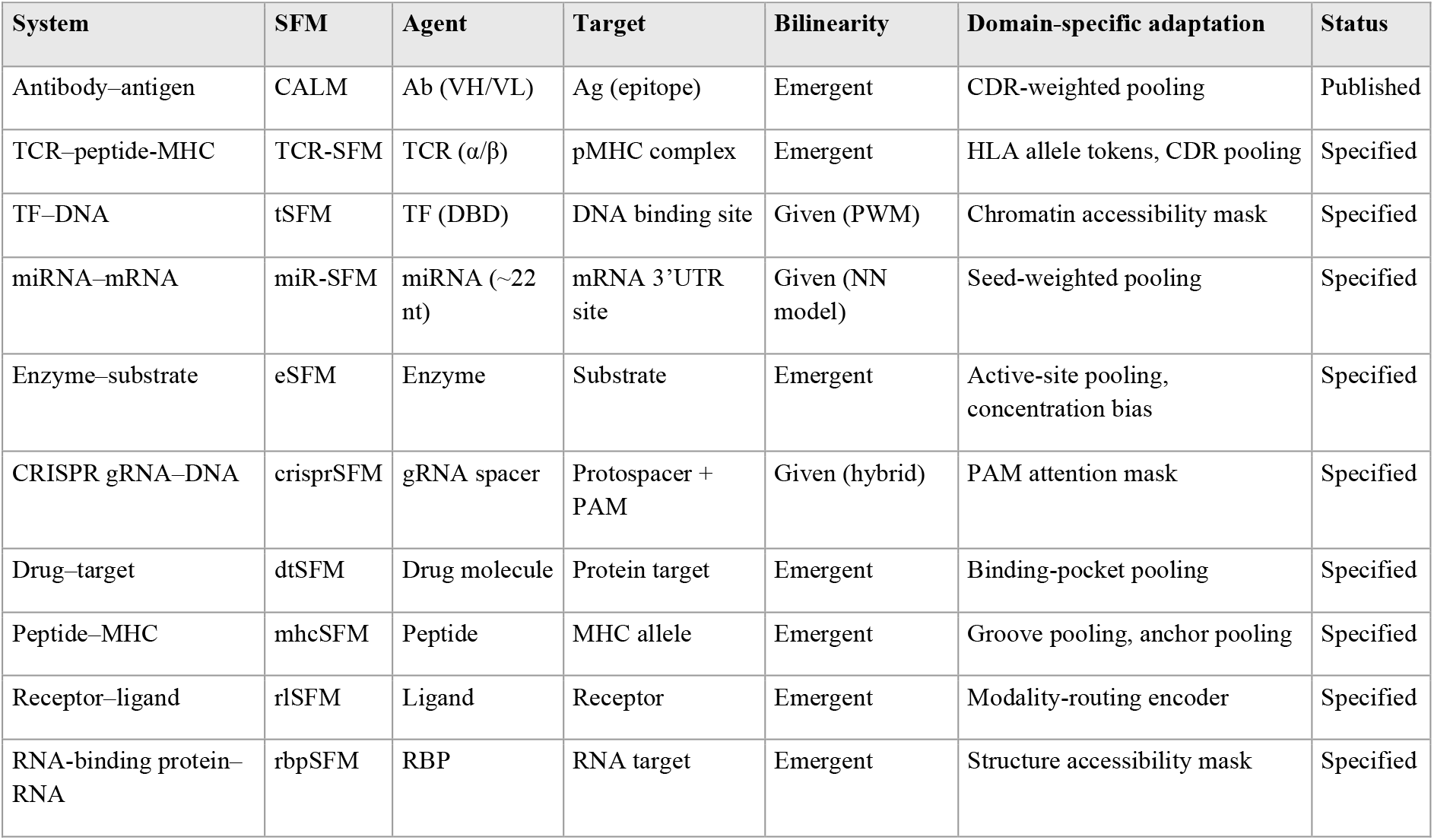
Ten molecular recognition systems verified against five architecture conditions.

All ten systems satisfy all five architecture conditions. The remaining conditions (discrete contacts, finite pools, thermal equilibrium, stochastic selection) are satisfied unconditionally by all ten; verification details are in Supplementary Note S2.

### 5.1 Domains with Published Results

#### Antibody–antigen

The Cross-attention Adaptive immune receptor-Antigen Language Model (CALM) represents the first SFM. Antibody CDR loops contact antigen epitope surfaces through 10– 30 residue-residue contact pairs (Chothia & Lesk, *J. Mol. Biol*., 1987). CALM uses AntiBERTy (Ruffolo *et al., Patterns*, 2022) and ESM-2 (Lin *et al., Science*, 2023) as dual encoders, trained on 4,138 pairs from SAbDab (Dunbar *et al., Nucleic Acids Res*., 2014). At 80% antigen identity clustering: R@1 6.5%/5.4% (Ag→Ab / Ab→Ag), R@10 17.0%/15.9%, against a random baseline of ∼0.2% (Table 2, Lee *et al., bioRxiv*, 2026).

### 5.2 Domains with Specified Architectures

The following nine domains have complete architectural specifications derived from the convergence equation and the five architecture conditions. Domain-specific data sources, curation protocols, and encoder configurations are detailed in Supplementary Table S1.

#### TCR–peptide-MHC (TCR-SFM)

Structurally homologous to antibody–antigen binding. TCR α/β chains use CDR loops to contact peptide residues presented in the MHC groove (Rudolph *et al., Annu. Rev. Immunol*., 2006). Bilinearity is emergent. The target is a two-component complex (peptide + MHC allele), requiring concatenated encoding with allele-specific tokens.

#### Transcription factor–DNA (tSFM)

The tightest mathematical correspondence in the program. Binding energy decomposes exactly by position through position weight matrices (Stormo, *Bioinformatics*, 2000), satisfying bilinearity in raw sequence space without learning. TF occupancy follows Boltzmann statistics at equilibrium (Bintu *et al., Curr. Opin. Genet. Dev*., 2005). Chromatin accessibility masking restricts the effective competitor pool to cell-type-specific open genomic regions, implemented as an attention mask.

#### MicroRNA–mRNA (miR-SFM)

Seed-region complementarity (positions 2–8) dominates the energy landscape (Bartel, *Cell*, 2009). Binding energy decomposes under the nearest-neighbor thermodynamic model (SantaLucia, *PNAS*, 1998). The cross-modal verification signal — computed duplex free energy *ΔG* via thermodynamic calculators — is the exact Boltzmann energy, making pseudo-labels for this domain theoretically tighter than structural proxy metrics used in protein domains.

#### Enzyme–substrate (eSFM)

At thermal equilibrium with equal substrate concentrations, enzyme– substrate selection follows Equation 2 directly. The Michaelis–Menten equilibrium assumption is standard enzymology (Michaelis & Menten, *Biochem. Z*., 1913). When substrate concentrations are unequal, a concentration-weighted bias ln[*S*_*j*_] is added to the attention logits — a biophysically motivated additive term. Active-site-weighted pooling restricts the agent encoding to catalytic residues.

#### CRISPR guide RNA–DNA (crisprSFM)

Guide RNA–target hybridization follows a position-weighted energy decomposition with a seed-to-distal mismatch tolerance gradient (Hsu *et al., Nat. Biotechnol*., 2013). PAM recognition functions as a mandatory attention mask: positions lacking a valid PAM are set to −∞ in the attention logits. The SFM models Phase 1 (reversible target selection), not Phase 2 (irreversible cleavage commitment) (Jinek *et al., Science*, 2012).

#### Drug–target selectivity (dtSFM)

Predicts which protein targets a drug molecule will bind across the proteome. Most approved drugs bind 5–30 protein targets; off-target binding drives approximately 30% of clinical trial failures. The bidirectional architecture enables both virtual screening (target → drug) and selectivity profiling (drug → target) in a single model. The drug encoder uses a molecular language model or graph neural network; the target encoder uses a protein language model with binding-pocket-weighted pooling.

#### Peptide–MHC presentation (mhcSFM)

Predicts which peptides will be presented by which MHC alleles — the central gating event in adaptive immunity. The human HLA locus encompasses over 13,000 class I alleles with extreme polymorphism concentrated in the peptide-binding groove. Existing methods (NetMHCpan, MHCflurry) achieve high accuracy on well-characterized alleles but do not learn a shared metric space enabling zero-shot prediction for rare alleles. The mhcSFM uses allele-groove-weighted pooling and anchor-position-weighted pooling.

#### Receptor–ligand signaling (rlSFM)

Cell surface receptors — approximately 800 GPCRs, 58 RTKs, 48 nuclear receptors, ∼40 cytokine receptors, and over 300 ion channels — mediate virtually all intercellular communication. The distinctive architectural challenge is ligand modality diversity: receptors bind small molecules, peptides, and proteins. The rlSFM uses a modality-routing encoder that handles all three ligand types in a unified framework.

#### RNA-binding protein–RNA (rbpSFM)

Approximately 1,500 human RNA-binding proteins control post-transcriptional gene expression. Each RBP recognizes specific RNA sequence motifs and/or structural elements. Existing methods are single-RBP models requiring retraining for each new protein. The rbpSFM learns a shared metric space across all RBPs, with RNA secondary structure accessibility masking analogous to chromatin accessibility masking in tSFM.

### 5.3 Scope and Extensibility

The ten domains above are those for which we have verified the five architecture conditions in detail and specified complete SFM architectures. They do not exhaust the scope of the convergence equation.

Any molecular recognition event at thermal equilibrium satisfying the five architecture conditions prescribes an SFM. Biological processes not included in the current program but governed by the same equation include: codon–anticodon selection during translation, DNA polymerase nucleotide selection during replication, chaperone–substrate recognition during protein folding, ubiquitin– substrate recognition during proteolytic targeting, histone modification reader–chromatin recognition during epigenetic regulation, spliceosome–splice site recognition during mRNA processing, olfactory receptor–odorant recognition during chemosensation, and neurotransmitter–receptor recognition during synaptic transmission. Each involves competitive molecular selection from a finite pool at thermal equilibrium. Each would converge to the same base architecture if an SFM were built for it. The ten specified domains represent the systems for which training data and domain expertise are currently available, not the theoretical boundary of the framework.

## §6. The MRC Scaling Law

### 6.1 Exponential Scaling from Architecture-Physics Alignment

Standard neural scaling laws describe test loss decreasing as a power law with dataset size: *L* ∝ *N*^−*α*^, with *α* typically 0.05–0.10 (Kaplan *et al., arXiv*, 2020; Hoffmann *et al., arXiv*, 2022). This regime characterizes function approximation in an unknown function class: the model must discover both the functional form and its parameters from data, and accuracy improves slowly with scale.

Level 3 architecture-physics alignment (§4) predicts a different regime. When the architecture implements the governing equation, the model does not learn how to convert scores into selection probabilities — the architecture does this by construction. It learns only the parameters that populate the known equation: the embedding vectors that determine compatibility scores between molecular agents and targets. This is parameter estimation in a known equation, not function approximation, and parameter estimation converges exponentially:

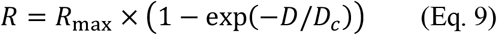

where *R* is retrieval accuracy, *R*_max_ is the asymptotic maximum achievable at infinite data, *D* is the effective diversity of training data (number of unique binding relationships), and *D*_*c*_ is a domain-specific constant we term the **diversity constant**. The exponential form predicts rapid improvement in the data-limited regime followed by saturation — qualitatively different from the slow, indefinite improvement of power-law scaling.

The critical variable is *D*, not *N*. Data diversity — the number of distinct binding relationships represented in the training set — drives learning, not data volume.

### 6.2 The Diversity Constant

The diversity constant *D*_*c*_ determines how much binding relationship diversity is required to saturate the exponential. It is predicted to correlate with thermodynamic degrees of freedom at the binding interface: systems with more contact positions, greater chemical diversity per position, and more conformational flexibility require more training data to learn the physical parameters.

This prediction is domain-specific:

**Given bilinearity domains** (tSFM, miR-SFM, crisprSFM): the energy decomposition is exact in raw sequence space, so the number of free parameters is determined by the number of positions and the alphabet size at each position. These systems have fewer effective degrees of freedom and are predicted to have smaller *D*_*c*_ — faster saturation, less training data required.

**Emergent bilinearity domains** (CALM, TCR-SFM, eSFM, dtSFM, mhcSFM, rlSFM, rbpSFM): the energy decomposition must be learned from data by the encoder, introducing additional parameters for the mapping from sequence to binding features. These systems have more effective degrees of freedom and are predicted to have larger *D*_*c*_ — slower saturation, more training data required.

If the correlation between *D*_*c*_ and thermodynamic degrees of freedom is observed across domains, it confirms that Level 3 alignment restricts the learning problem to physical parameters: the amount of data required to learn the model is determined by the physical complexity of the binding interface, not by the capacity of the neural network.

### 6.3 Model Selection

For each domain, both exponential (Eq. 9) and power-law fits are compared using the Bayesian Information Criterion (Kass & Raftery, *Journal of the American Statistical Association*, 1995). The protocol uses diversity-controlled training subsets at 10%, 25%, 50%, 75%, and 100% of maximum binding pair diversity, with volume-controlled subsets at fixed diversity to deconfound diversity from volume. *Δ*BIC > 10 constitutes decisive evidence for the preferred model.

The prediction is that *Δ*BIC > 10 (exponential preferred over power-law) in all domains. If power-law scaling is observed instead in any domain, the Level 3 alignment claim for that domain is undermined: it would indicate that the architecture is not fully implementing the governing equation for that system, and that the model is performing function approximation rather than parameter estimation. Power-law scaling in a specific domain is therefore a diagnostic, not a failure of the general theory — it indicates that one or more of the five architecture conditions is not adequately satisfied in the current implementation.

### 6.4 Relationship to Shannon’s Channel Capacity

Shannon proved that the theoretical limit of a communication channel — the maximum rate at which information can be transmitted reliably — is determined by physical properties of the channel (bandwidth, signal-to-noise ratio) and cannot be exceeded by any encoding scheme (Shannon, *Bell System Technical Journal*, 1948). The MRC scaling law predicts an analogous limit for molecular specificity prediction: the maximum achievable retrieval accuracy for a given amount of training data diversity, determined by binding interface physics (*D*_*c*_). The matched filter theorem (North, *RCA Technical Report*, 1943) provides a further parallel: once the optimal filter is known (the convergence equation settles the architecture), what remains is signal processing (data engineering).

These parallels are, at this stage, aspirational rather than demonstrated. Shannon’s channel capacity is a proven theorem. The MRC scaling law is a prediction. The distinction between a theorem and a prediction is precisely the gap between the mathematical framework (established) and its empirical consequences (untested). Closing this gap requires training and evaluating SFMs across multiple domains at diversity-controlled data scales.

## §7. Affinity Calibration

The SFM architecture as described in §3 is a discriminative system: it predicts *which* target an agent binds (the retrieval problem). This section derives from the convergence equation that the same architecture also predicts how tightly the agent binds (the affinity problem), with only two additional parameters.

### 7.1 Two-Parameter Sufficiency

The convergence equation identifies the contrastive similarity score with the Boltzmann energy:

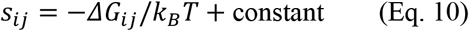

The additive constant is the thermodynamic reference state. It cancels in the softmax denominator, which is why contrastive training recovers correct rankings without ever observing a measured binding affinity. But the constant remains in the absolute score. To recover quantitative binding free energies from contrastive scores, the constant must be calibrated out.

A two-parameter linear regression is sufficient:

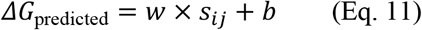

where the slope *w* absorbs the unit conversion from embedding-space units to kcal/mol (including any scaling by *τ*), and the intercept *b* absorbs the thermodynamic reference state (the baseline free energy of the unbound system, including solvation entropy and standard-state concentration conventions).

The two-parameter sufficiency is a direct consequence of the convergence equation. If the identity *s*_*ij*_ = −*ΔG*_*ij*_/*k*_*B*_*T* + constant holds quantitatively, then the relationship between contrastive score and binding free energy is linear, and no higher-order terms are needed. The sufficiency is itself testable: if a quadratic or higher-order model is significantly preferred by BIC over the linear model, it indicates a systematic nonlinearity in the embedding space that the convergence equation does not account for (Guo *et al., ICML*, 2017).

### 7.2 Physical Interpretation of the Regression Parameters

The slope *w* encodes the temperature and unit conversion. In the Boltzmann identity, *s*_*ij*_ = −*ΔG*_*ij*_/*k*_*B*_*T* implies *ΔG*_*ij*_ = −*k*_*B*_*T* × *s*_*ij*_. If the embedding space were perfectly calibrated in energy units, *w* = −*k*_*B*_*T* = −0.616 kcal/mol at 310 K. In practice, the embedding-space dot product has an arbitrary scale determined by the projection head dimension, L2 normalization, and the learned *τ*, so *w* absorbs these factors. The sign of *w* should be negative (higher similarity corresponds to more negative *ΔG*, i.e., tighter binding).

The intercept *b* is the thermodynamic reference state: the free energy of the unbound system, including solvation, configurational entropy, and standard-state conventions. None of these quantities are represented in the contrastive embedding; the intercept anchors the embedding-space scores to the experimental reference frame.

A specific prediction follows: if the regression is performed on both post-*τ* scores (*s* = ⟨*z*_agent_, *z*_target_⟩/*τ*) and pre-*τ* raw dot products (*s*′ = ⟨*z*_agent_, *z*_target_⟩), the ratio of slopes should equal the learned temperature: *w*_*B*_/*w*_*A*_ = *τ*_learned_. This is the **temperature recovery** prediction (§3.6): if the ratio holds, the learned *τ* has the physical meaning assigned by the convergence equation.

### 7.3 The Decoupling Principle

The affinity calibration framework reveals a fundamental asymmetry in the data requirements for specificity prediction.

The contrastive training data — binary binding pairs indicating that agent *i* binds target *j* — teaches the geometry of the embedding space: the relative positions and orientations of all agents and targets such that binding partners are close and non-partners are far. Under the convergence equation, this geometry is the energy landscape. Each dot product ⟨*z*_agent_, *z*_target_⟩ is a relative binding energy. Scaling from thousands to millions of contrastive pairs refines this geometry across a larger and more diverse region of molecular space.

The calibration data — measured *K*_*d*_ values converted to *ΔG* = *RT*ln(*K*_*d*_) — teaches two numbers: the slope and the intercept. These are properties of the coordinate system, not of the data distribution. They do not change as the landscape becomes more detailed.

This is **the decoupling principle**: the energy landscape scales with cheap data (binary binding pairs), and the unit conversion is a fixed-cost operation requiring a small number of expensive measurements (*K*_*d*_ values from SPR, ITC, or BLI). The **leverage ratio** — the ratio of cheap binding pairs to expensive affinity measurements required — quantifies this asymmetry. The leverage ratio grows with contrastive data scale: at 4,000 contrastive pairs with 500 calibration points, the ratio is approximately 8:1; at one million contrastive pairs with the same 500 calibration points, the ratio is 2,000:1.

The decoupling principle applies to every SFM domain where affinity data is scarce but binding data is abundant. Domains with the highest leverage ratio are those where the SFM approach most dramatically outperforms conventional affinity-first methods. Full cross-domain leverage ratio predictions are provided in Supplementary Note S4.

### 7.4 Cross-Domain Predictions

The affinity calibration framework makes domain-specific predictions that connect to the bilinearity classification (§3.1, §5).

For **given bilinearity** domains (tSFM, miR-SFM, crisprSFM), the embedding space is structurally closer to the true energy function because the energy decomposition is exact in raw sequence space. The slope *w* should converge closer to the physical prediction (−*k*_*B*_*T* up to a scaling factor) and the linearity test should show negligible higher-order terms.

For **emergent bilinearity** domains (CALM, TCR-SFM, eSFM, dtSFM, mhcSFM, rlSFM, rbpSFM), the embedding space must learn the energy function from data, introducing additional degrees of freedom. The slope *w* has more freedom, and systematic nonlinearities are more likely — particularly at the extremes of the affinity range, where L2 normalization may compress the high-affinity tail (cosine similarity saturates at 1.0).

If this pattern is observed across domains — tighter linearity and more physical slope values in given-bilinearity domains, greater nonlinearity in emergent-bilinearity domains — it directly connects the affinity calibration results to the bilinearity classification and provides additional evidence for the physical interpretation of the embedding space.

## §8. Hybrid Recursive Learning

Sections 3–7 describe the discriminative and generative SFM: an architecture that retrieves binding partners and generates novel binding sequences from sequence alone. This section derives a theoretical framework for improving SFM performance through reinforcement learning with synthetic verification, without requiring experimental labels in the optimization loop.

### 8.1 The Experimental Constraint

Training data for molecular recognition models is generated by experimental platforms that impose a fundamental asymmetry: they can vary one binding partner while holding the other fixed, but cannot co-vary both simultaneously at scale.

Phage display (Smith, *Science*, 1985; McCafferty *et al., Nature*, 1990) and yeast surface display (Boder & Wittrup, *Nat. Biotechnol*., 1997) construct libraries of 10^7^ – 10^11^ protein variants and select for binding to a fixed target. Deep mutational scanning (Fowler & Fields, *Nat. Methods*, 2014) and deep mutational learning (Mason *et al., Nat. Biomed. Eng*., 2021; Taft *et al., Cell*, 2022) extend this by coupling display selection with deep sequencing and machine learning, enabling sequence-to-function prediction within the neighborhood of a single antibody or antigen. But the training data is inherently one-sided: antibody variants are screened against a fixed antigen, or antigen variants are screened against a fixed antibody. The joint space — novel antibodies against novel antigens, novel enzymes against novel substrates, novel TCRs against novel peptide-MHC complexes — cannot be explored experimentally at scale because it requires constructing and screening combinatorial libraries in both binding partners simultaneously.

This is the constraint that hybrid recursive learning addresses. The SFM decoder (§3.7) proposes candidates in the joint space computationally, and a structural verifier evaluates them without experimental assays. The optimization loop operates entirely in silico, enabling exploration of binding partner combinations that no experimental platform can access.

### 8.2 The Actor-Verifier Decomposition

**Hybrid recursive learning** decomposes the optimization into two components operating in different modalities.

#### Actor

The SFM decoder proposes candidate sequences conditioned on the target embedding, operating in sequence space. Two decoder architectures are prescribed: autoregressive (AR) decoding, which generates tokens sequentially (Raffel *et al., JMLR*, 2020), and discrete diffusion (DD) decoding, which generates all positions in parallel through iterative denoising (Austin *et al., NeurIPS*, 2021; Sahoo *et al., arXiv*, 2024). The DD decoder is the more physics-aligned choice: in the Boltzmann framework, binding occupancy at each target position is determined simultaneously at equilibrium, not sequentially. The autoregressive factorization *P*(*x*_1_)*P*(*x*_2_|*x*_1_) ⋯ *P*(*x*_*L*_|*x*_<*L*_) imposes an artificial left-to-right ordering that the binding physics does not have. Both architectures share the same cross-attention interface to the dual encoders (§3.7); the choice between them is empirical and domain-dependent.

#### Verifier

A structure-prediction or thermodynamic model evaluates each candidate in a representation space orthogonal to the actor’s token sequence. The class of available verifiers spans multiple modalities:

For protein domains, co-folding models predict the 3D structure of the agent-target complex and return interface metrics. The current generation includes AlphaFold-Multimer (Evans *et al., bioRxiv*, 2022), AlphaFold3 (Abramson *et al., Nature*, 2024), RoseTTAFold All-Atom (Krishna *et al., Science*, 2024), Boltz-1 and Boltz-2 (Wohlwend *et al*., 2024, 2025), and Chai-1 (Chai Discovery, 2024). These models generate structural ensembles from which calibrated binding probabilities, interface metrics (buried surface area, hydrogen bond counts, clash scores), and confidence estimates can be derived. The structure-based protein design models RFdiffusion (Watson *et al., Nature*, 2023) and Proteina-Complexa (Didi *et al., ICLR*, 2026) provide additional structural evaluation capabilities. For protein-ligand interactions, molecular docking tools including AutoDock Vina (Trott & Olson, *J. Comput. Chem*., 2010), GNINA (McNutt *et al., J. Cheminform*., 2021), and DiffDock (Corso *et al., ICLR*, 2023) evaluate binding poses and estimate binding free energies.

For nucleic acid domains, thermodynamic calculators compute duplex or secondary structure free energies directly from sequence. ViennaRNA (Lorenz *et al., Algorithms Mol. Biol*., 2011) and NUPACK (Zadeh *et al., J. Comput. Chem*., 2011) compute *ΔG* for RNA-RNA, DNA-RNA, and DNA-DNA duplexes using experimentally parameterized nearest-neighbor models (SantaLucia, *PNAS*, 1998). These verifiers compute the exact Boltzmann energy — the verification signal is not a proxy but the physical quantity itself.

All verifiers share the property that they operate in a representation space (3D coordinates or thermodynamic energies) that is orthogonal to the actor’s token-sequence space. This orthogonality is the theoretical basis for the framework’s robustness (§8.3).

The actor and verifier implement the convergence equation through fundamentally different mathematical machinery. The actor learns −Δ*G*/*k*_B_*T* through contrastive co-embedding and AR/DD decoding over token sequences. The verifier estimates *ΔG* through geometric interface energy computation or thermodynamic calculation. They model the same physical quantity — binding free energy — through orthogonal representations.

The optimization follows group relative policy optimization (GRPO): for each target, the actor proposes a group of *G* candidates; the verifier scores each; the policy update uses within-group relative advantages with a KL divergence penalty to a frozen reference policy (DeepSeek-AI, *arXiv*, 2025). No learned critic or value network is required. This approach was demonstrated at scale for language model reasoning by DeepSeek-R1, which showed that verifier-only RL with GRPO is stable without learned reward models or human raters, provided the verifier is reliable and training is regularized (DeepSeek-AI, *arXiv*, 2025). The standard RLHF framework (Ouyang *et al., NeurIPS*, 2022; Schulman *et al., arXiv*, 2017) is a special case where the verifier is a learned reward model trained on human preferences; hybrid recursive learning replaces this with a physics-based verifier.

### 8.3 The Orthogonal Verification Principle

The theoretical basis for hybrid recursive learning is that verification across orthogonal inductive biases is more robust than verification within a single modality.

In same-modality verification, the actor and verifier share representational structure. Systematic biases in the shared representation propagate to both the proposal and the evaluation. Reward hacking exploits these shared biases: the actor learns to generate candidates that score well according to the shared bias, not according to the physical truth.

In cross-modal verification, the actor and verifier have orthogonal inductive biases. The sequence-space actor (autoregressive transformer; Vaswani *et al., NeurIPS*, 2017) and the structure-space verifier (SE(3)-equivariant diffusion model; Fuchs *et al., NeurIPS*, 2020) share no architectural components, no training data, no loss functions, and no representational primitives. Their error modes are decorrelated: a candidate that exploits a bias in the sequence model’s token-level likelihood is unlikely to simultaneously exploit a bias in the structure model’s geometric energy function, because the two representations have no structural overlap.

Formally: let *ϵ*_actor_ and *ϵ*_verifier_ denote the systematic error distributions of the actor and verifier, respectively. In same-modality verification, Corr(*ϵ*_actor_, *ϵ*_verifier_) is high because both models share representational structure. In cross-modal verification, the orthogonality of inductive biases predicts Corr(*ϵ*_actor_, *ϵ*_verifier_) ≈ 0. Agreement between models with uncorrelated errors is evidence of physical truth; agreement between models with correlated errors may be evidence of shared bias. This is the **orthogonal verification principle**: the reliability of synthetic verification scales with the decorrelation of errors between actor and verifier.

### 8.4 Collapse Resistance

Policy collapse — where the actor converges to a narrow distribution of high-reward but non-diverse outputs — is the primary failure mode of reinforcement learning with synthetic rewards (Amodei *et al., arXiv*, 2016). The orthogonal verification principle provides a theoretical basis for collapse resistance in hybrid recursive learning, supplemented by established regularization techniques.

#### Error decorrelation

Cross-modal orthogonality raises the bar for reward hacking. To exploit the reward signal, the actor must generate candidates that simultaneously fool two fundamentally different mathematical frameworks — decoder-generated token likelihood and SE(3)-equivariant geometric energy. This is strictly harder than fooling a single-modality verifier.

#### KL regularization

A KL divergence penalty between the current policy π_θ_ and a frozen reference policy π_ref_ preserves the base competencies of the actor — fluency, diversity, and physical plausibility of generated sequences. This is standard in RLHF (Ouyang *et al., NeurIPS*, 2022) and was shown to be sufficient for stability in verifier-only RL (DeepSeek-AI, *arXiv*, 2025).

#### Group-relative optimization

GRPO computes advantages relative to the group mean: for each target, a group of *G* candidates is proposed and scored, and the advantage of each candidate is its reward minus the group median. This eliminates the learned critic — a common source of instability in actor-critic methods (Schulman *et al., arXiv*, 2017) — and reduces variance through within-group normalization.

#### Verifier abstention

When the structural verifier’s ensemble agreement falls below a confidence threshold, it abstains — returning a reward of zero rather than a noisy estimate. This prevents the actor from learning on unreliable signals and bounds the noise in the reward surface. Calibrated abstention is essential: an uncalibrated verifier that always returns a score, regardless of confidence, provides a reward surface that the actor can exploit (Guo *et al., ICML*, 2017).

The combination of orthogonal verification, KL regularization, GRPO, and verifier abstention provides four independent collapse resistance mechanisms operating at different levels: representational (orthogonality), distributional (KL), optimization (GRPO), and signal quality (abstention).

### 8.5 Generalization Across SFM Domains

The hybrid recursive learning framework generalizes beyond any single domain. For each SFM, the actor is the domain-specific decoder (§3.7) and the verifier is the appropriate structural or thermodynamic

| SFM domain | Actor (sequence space) | Verifier (structure/energy space) |
| --- | --- | --- |
| CALM, TCR-SFM | SFM decoder | AlphaFold3, Boltz-2, Chai-1, RoseTTAFold All-Atom |
| tSFM | SFM decoder | ViennaRNA (protein-DNA thermodynamics), JASPAR PWM energies |
| miR-SFM, crisprSFM | SFM decoder | ViennaRNA, NUPACK (duplex $\Delta G$) |
| eSFM | SFM decoder | AutoDock Vina, GNINA, DiffDock |
| dtSFM | SFM decoder | AutoDock Vina, GNINA, DiffDock, FEP+ |
| mhcSFM | SFM decoder | AlphaFold3, NetMHCpan (structure-based) |
| rlSFM, rbpSFM | SFM decoder | AlphaFold3, Boltz-2, domain-specific structural models |

For nucleic acid domains (tSFM, miR-SFM, crisprSFM), the verifier computes duplex free energy *ΔG* directly — the exact Boltzmann energy (SantaLucia, *PNAS*, 1998). The verification signal in these domains is theoretically tighter than in protein domains, where the verifier estimates *ΔG* through structural proxy metrics such as interface ipTM or DockQ scores. For protein-ligand domains (eSFM, dtSFM), molecular docking provides pose-level evaluation, and free energy perturbation methods (FEP+; Wang *et al., J. Am. Chem. Soc*., 2015) provide higher-accuracy but computationally expensive binding free energy estimates. The orthogonal verification principle holds in all cases: the actor operates on token sequences and the verifier operates on energies or geometries.

### 8.6 Scope

This section prescribes the hybrid recursive learning framework from theory and derives the orthogonal verification principle as the theoretical basis for collapse resistance. The implementation —specific verifier architectures, GRPO hyperparameters, KL schedules, abstention thresholds, reward surface design — is domain-dependent engineering. The theoretical claim is that cross-modal actor-verifier decomposition with orthogonal inductive biases provides a principled foundation for reinforcement learning with synthetic rewards in molecular design, and that the convergence equation —governing both the actor’s sequence-space objective and the verifier’s structure-space evaluation— ensures that agreement between modalities is evidence of physical truth rather than shared computational artifact.

## §9. Sequence-First Versus Structure-First Paradigms

### 9.1 Two Paradigms

The dominant paradigm in computational molecular biology assumes that the path from sequence to function passes through structure: Sequence → Structure → Function. This paradigm encompasses structure prediction models — AlphaFold2 (Jumper *et al., Nature*, 2021), AlphaFold3 (Abramson *et al., Nature*, 2024), RoseTTAFold All-Atom (Krishna *et al., Science*, 2024), Boltz-1/Boltz-2 (Wohlwend *et al*., 2024, 2025), Chai-1 (Chai Discovery, 2024) — structure-based design models — RFdiffusion (Watson *et al., Nature*, 2023), ProteinMPNN (Dauparas *et al., Science*, 2022), BoltzGen (Stark *et al., bioRxiv*, 2025) — and gradient-based binder optimization methods — BindCraft (Pacesa *et al., Nature*, 2025), which backpropagates through AlphaFold2-Multimer’s predicted interface confidence (ipTM) to directly optimize binder sequences against a fixed target structure, and Proteina-Complexa (Didi *et al., ICLR*, 2026). We term this the **structure-first paradigm**.

The framework developed in this paper follows a different path: Sequence → Specificity, governed by the convergence equation. We term this the **sequence-first paradigm**. The distinction is mathematical, and the mathematical differences have consequences for learnability, data efficiency, and computational cost.

### 9.5 Mapping Cardinality and Problem Formulation

Every supervised learning problem defines a mapping from inputs to outputs. The learnability of the mapping depends fundamentally on its cardinality (Vapnik, *The Nature of Statistical Learning Theory*, 1995).

**Structure prediction (Sequence → Structure)** is a many-to-one mapping: each sequence folds into one structure (Anfinsen, *Science*, 1973). This is a well-posed learning problem — there is a unique correct answer for each input, standard supervised learning applies, and AlphaFold2 solved it decisively (Jumper *et al., Nature*, 2021).

**Inverse folding (Structure → Sequence)** is a one-to-many mapping: for any given protein fold, thousands to millions of amino acid sequences are compatible. Proteins sharing less than 20% sequence identity routinely adopt identical folds (Rost, *Protein Engineering*, 1999); the TIM barrel fold alone encompasses thousands of unrelated sequences (Nagano *et al., J. Mol. Biol*., 2002). Nature contains approximately 1,500 distinct fold families (Chothia, *Nature*, 1992; Orengo *et al., Structure*, 1997), while the sequence space for a 100-residue protein is approximately 20^100^. This is an ill-posed problem: there is no unique correct answer, the model is sampling from a vast distribution, and training data (the PDB) provides usually only one example sequence per structure out of thousands of valid alternatives. Models trained on this data — including ProteinMPNN (Dauparas *et al., Science*, 2022) — optimize sequence recovery (reproducing the native sequence) rather than functional fitness (producing sequences that bind a target), because native sequence recovery is the available supervision signal.

**Specificity prediction (Sequence → Binding probability)** is a many-to-one mapping: many sequence pairs map to the same binding probability. This is a well-posed learning problem with a unique correct answer (the Boltzmann probability) for each input. The convergence equation specifies both the functional form and the training objective (InfoNCE, §3.3).

The structure-first approach to specificity prediction chains a well-posed problem (Sequence → Structure) with an ill-posed problem (Structure → Sequence for binding partners), followed by a scoring step (Structure → Binding estimate). The ill-posed intermediate step — generating candidate binding sequences from structure — is the mathematical bottleneck. The sequence-first approach bypasses this step entirely: Sequence → Binding probability, many-to-one, well-posed.

### 9.3 Information Flow and Dimensionality

The structure-first specificity pipeline follows the information flow: Structure (∼10^3^ fold families, low-dimensional) → Sequence (∼20^100^ possible sequences, high-dimensional) → Binding affinity (scalar, low-dimensional). The middle step expands from a low-dimensional representation into an astronomically larger space, and the final output collapses back to a scalar. This Low → High → Low trajectory introduces an intermediate expansion that creates a vast space of candidates, most of which are irrelevant to the final output.

The sequence-first pipeline follows the information flow: Sequence pair (high-dimensional, two molecular sequences) → Binding probability (scalar, low-dimensional). This is monotonic compression — High → Low — with no intermediate expansion. The dual encoders compress each sequence into an embedding vector; the dot product compresses the pair into a scalar score; the convergence equation converts the score into a probability. At each stage, dimensionality decreases. No information is invented; features are selected and noise is discarded.

From an information-theoretic perspective, the structure-first intermediate expansion moves in the direction of increasing information entropy: generating a high-dimensional sequence from a low-dimensional structural specification requires the model to invent information not present in the input. The sequence-first compression moves in the direction of decreasing information entropy: extracting a binding probability from sequences requires the model to select relevant features and discard irrelevant variation. Compression is a well-characterized operation in statistical learning; expansion requires generating information not determined by the input, which is a harder problem in general (Shannon, *Bell System Technical Journal*, 1948).

### 9.4 Loss Function Alignment

The structure-first pipeline trains its components on proxy objectives that diverge from the specificity prediction goal.

ProteinMPNN optimizes sequence recovery: given a backbone structure, reproduce the native amino acid sequence (Dauparas *et al., Science*, 2022). But sequence recovery and binding affinity are different objectives — many non-native sequences would bind the target; many sequences similar to the native would not. The loss function penalizes deviation from the native sequence, which means it penalizes valid solutions that differ from the single observed natural sequence.

AlphaFold2 and AlphaFold3 optimize structural accuracy (Fram Aligned Point Error or FAPE, diffusion loss over atomic coordinates). These objectives do not include any term for binding specificity (Abramson *et al., Nature*, 2024). Interface confidence metrics such as ipTM and DockQ (Mariani *et al., Bioinformatics*, 2013) measure structural plausibility, not thermodynamic favorability —a distinction that has been shown to limit their utility as binding affinity predictors. In one of the largest systematic evaluations of computational protein design metrics, the Adaptyv EGFR binder design competition selected and characterized 601 designed proteins for expression and binding affinity to EGFR using automated bio-layer interferometry. Across all structure prediction confidence metrics evaluated — including AlphaFold2’s ipTM, iPAE, and ESM2 pseudo-log-likelihood — no significant correlation with experimentally measured binding affinity was found, and for most metrics the observed trend was opposite to the expected direction: designs with higher structural confidence scores did not exhibit stronger binding (Cotet *et al., bioRxiv*, 2025).

The sequence-first paradigm optimizes the convergence equation directly via InfoNCE (§3.3). The training objective IS the physical selection probability.

### 9.5 Training Distribution and Out-of-Distribution Failure

A further mathematical difficulty for the structure-first paradigm arises from training distribution mismatch. AlphaFold2, AlphaFold3, and related co-folding models are trained on experimentally determined structures from the PDB — natural proteins shaped by evolutionary selection. Computationally designed protein sequences, particularly those generated by diffusion-based backbone generation followed by inverse folding, occupy a different region of sequence space than natural proteins. They may satisfy structural constraints (correct fold) while violating the statistical regularities of natural protein sequences on which the structural models were trained.

This creates a compounding error: the design pipeline generates sequences that are out-of-distribution for the validation model (AlphaFold2), which then serves as a filter. The filter’s reliability degrades precisely on the sequences it is asked to evaluate — an adversarial dynamic that worsens as designs become more novel. Empirical evidence supports this concern: co-folding models have been shown to produce physically unrealistic structures when evaluated on designed sequences that deviate from natural sequence statistics, suggesting that the models learn statistical patterns in their training data rather than the underlying physics of binding.

SFMs are not immune to distribution shift, but the nature of the shift is different. The SFM is trained on binding pairs (agent-target relationships), and generalization is measured by performance on novel agents, novel targets, or novel pairs not seen during training. The leakage-controlled evaluation framework used in CALM — clustering by antigen identity at 40%, 60%, and 80% thresholds (Lee *et al., bioRxiv*, 2026) — explicitly measures this out-of-distribution performance. The distribution shift is in the binding relationship space, not in the sequence statistics space.

### 9.6 Gradient-Based Design and Unified Design-Prediction Models

Two recent approaches within the structure-first paradigm merit specific discussion because they address some of the limitations identified above.

#### Gradient-based binder optimization

BindCraft (Pacesa *et al., Nature*, 2025) backpropagates through the predicted interface confidence score (ipTM) of AlphaFold2-Multimer (Evans *et al., bioRxiv*, 2022) to directly optimize binder sequences against a fixed target structure. Rather than generating backbone structures and then designing sequences (the RFdiffusion → ProteinMPNN pipeline), BindCraft optimizes sequences end-to-end through the structure prediction model’s differentiable computational graph. This eliminates the explicit backbone generation step and allows sequence optimization to be guided by the full structural model. However, the approach inherits two constraints. First, the optimization target remains ipTM — a structural confidence metric, not a thermodynamic quantity — so the loss function alignment limitation (§9.4) persists. Second, each optimization trajectory requires hundreds to thousands of forward and backward passes through AlphaFold2-Multimer, with each pass requiring minutes of GPU time, making the computational cost comparable to RFdiffusion-based pipelines (hundreds to thousands of GPU-hours per target). Third, the training distribution concern (§9.5) applies: the sequences discovered by gradient-based optimization may exploit features of the AlphaFold2 loss landscape that do not correspond to physical binding, particularly for sequences far from the natural protein distribution.

#### Unified design-prediction models

BoltzGen (Stark *et al., bioRxiv*, 2025) represents a more recent advance: a single all-atom diffusion model that jointly generates binder structure and sequence while simultaneously performing structure prediction. By unifying design and folding within one model using a geometric encoding of residue types, BoltzGen avoids the modular pipeline of separate generation, inverse folding, and validation steps. Experimental validation across 26 targets showed nanomolar binders for 66% of novel targets with less than 30% sequence similarity to any known bound structure. BoltzGen addresses several limitations of the modular pipeline — the cascade of correction steps is collapsed into a single model, and the geometric residue encoding enables joint optimization of structure and sequence. The mapping cardinality concern (§9.2) is partially mitigated because design and folding are unified rather than sequential. However, the model operates entirely in structure space: it generates 3D coordinates and derives sequence from geometry. The information flow remains structure-centric, and the optimization target is structural plausibility rather than the convergence equation. Whether unified structure-based approaches can solve molecular recognition —predicting *which* targets a designed binder will bind from among all possible targets, rather than designing a binder for a specified target — remains an open question that defines the boundary between the paradigms.

### 9.7 Precedent from Sequence-Based Specificity Models

The hypothesis that molecular sequence contains sufficient information for specificity prediction did not originate with the convergence framework. Experimental evidence from display technology and deep mutational learning demonstrated that narrow machine learning models trained on sequence data alone could accurately predict binding specificity within defined molecular neighborhoods.

Mason *et al*. showed that deep neural network classifiers trained on mammalian display screening data could predict antibody-antigen specificity from antibody sequence with approximately 85% precision for trastuzumab variants binding HER2 (Mason *et al., Nat. Biomed. Eng*., 2021). Taft *et al*. extended this to the antigen side, demonstrating that models trained on combinatorial mutagenesis libraries of the SARS-CoV-2 receptor-binding domain could predict ACE2 binding and antibody escape from sequence (Taft *et al., Cell*, 2022). These models were narrow — each was trained on variants of a single antibody or antigen and could not generalize beyond its training domain — but they established a critical principle: sequence holds sufficient information for specificity prediction, at least within defined molecular neighborhoods.

The convergence framework provides the theoretical explanation for why this works (the convergence equation operates on sequence-derived compatibility scores) and the architectural prescription for generalizing it across molecular space (dual encoders with contrastive training). The narrow models of Mason *et al*. and Taft *et al*. learned local regions of the binding energy landscape; the SFM architecture learns the global landscape by implementing the governing equation.

### 9.8 Computational Economics

The mathematical differences have direct consequences for computational cost. AlphaFold3-scale co-folding requires minutes to hours per complex on GPU hardware; evaluating one antibody against 10,000 candidate antigens requires 10,000 independent co-folding runs. RFdiffusion-based binder design requires hundreds to thousands of backbone generation trajectories per target, each followed by ProteinMPNN sequence design and AlphaFold2 structural validation — a pipeline of thousands of GPU-hours per target (Watson *et al., Nature*, 2023; Bennett *et al., Nature*, 2025).

An SFM evaluates the same ranking in a single forward pass through dual encoders followed by a matrix multiplication of the resulting embeddings — seconds on a single GPU. Training CALM required approximately 100–200 GPU-hours on A100 hardware for ∼4,000 pairs (Lee *et al., bioRxiv*, 2026). The minimum viable compute for useful specificity prediction differs by orders of magnitude between the paradigms — a direct consequence of the Level 3 alignment argument (§4) that the SFM architecture encodes the physics, reducing the learning problem to parameter estimation.

### 9.9 The Complementary Relationship

The structure-first paradigm has produced major advances in protein folding (Jumper *et al., Nature*, 2021), de novo protein design (Kuhlman *et al., Science*, 2003; Huang *et al., Nature*, 2016; Cao *et al., Science*, 2020), inverse folding (Dauparas *et al., Science*, 2022), and structure-based binder design (Watson *et al., Nature*, 2023; Stark *et al., bioRxiv*, 2025). The mathematical analysis above identifies specific limitations of the paradigm for molecular recognition — the specificity prediction problem — but does not diminish its contributions to the folding and design problems, which are distinct.

The two paradigms are complementary. Structural co-folding models serve as pseudo-labeling oracles for SFM training: they generate candidate molecular complexes, interface metrics filter for high-confidence pairs, and filtered pairs become contrastive training data for the sequence-based recognition model. For nucleic acid domains, thermodynamic calculators (ViennaRNA; Lorenz *et al., Algorithms Mol. Biol*., 2011; NUPACK; Zadeh *et al., J. Comput. Chem*., 2011) compute the exact Boltzmann energy as pseudo-labels. For protein-ligand domains, molecular docking programs provide pseudo-labels. Each verification tool generates labeled training data; the SFM learns specificity from that data.

At inference, SFMs with trained decoders (§3.7) are complete generative systems: they retrieve binding partners and generate novel binding sequences from sequence alone, without structural input. Structural models serve as training data infrastructure and are not required at deployment.

### 9.10 Falsification Framework

Both paradigms make testable predictions.

#### The sequence-first paradigm predicts

SFMs exhibit exponential scaling of retrieval accuracy with training data diversity (ΔBIC > 10 over power-law in all domains). The diversity constant *D*_*C*_ correlates with thermodynamic degrees of freedom. SFMs outperform structure-first scoring pipelines at matched compute budgets. Affinity calibration yields a linear relationship between contrastive scores and measured *ΔG* with two-parameter sufficiency.

#### The structure-first paradigm predicts

Continued improvements in co-folding accuracy and unified design-prediction models will produce reliable binding affinity predictions from structure. Structure-based approaches, with sufficient compute and structural data, will match or exceed sequence-first models for specificity prediction.

Neither set of predictions has been tested at sufficient scale. Both paradigms would benefit from controlled comparisons at matched compute budgets across multiple molecular recognition domains.

## §10. The Logical Structure of Failure

The preceding sections derive the convergence equation (§2), the architecture it prescribes (§3), the alignment classification (§4), ten domain-specific instantiations (§5), scaling predictions (§6), affinity calibration theory (§7), hybrid recursive learning (§8), and the paradigm comparison (§9). These are theoretical contributions. The empirical program — training SFMs across domains, measuring scaling behavior, validating predictions experimentally — is largely ahead.

The practical failure modes — insufficient training data diversity, systematic pseudo-label noise, and poor bilinearity approximation in specific protein domains — are analyzed in §10.2. The present section addresses the prior question of whether the framework can fail in principle, independent of data quality.

### 10.1 Two Conditions for Failure

Assuming sufficient training data of adequate quality — labeled binding pairs spanning the diversity of each molecular recognition domain — a Specificity Foundation Model can fail to learn molecular specificity only if one of two conditions holds.

**Condition 1: The convergence equation does not govern molecular binding at thermal equilibrium**. The framework rests on the claim that binding probability follows:

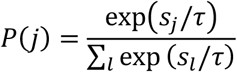

This is a consequence of the second law of thermodynamics. The Boltzmann distribution has been verified experimentally across every domain of chemistry and biophysics for over a century. Every *K*_*d*_ measurement made by surface plasmon resonance, isothermal titration calorimetry, or any other binding assay is a measurement of the Boltzmann probability ratio between bound and unbound states. The Luce uniqueness theorem (§2) proves that this is the only function satisfying the five axioms of competitive selection — any alternative violates at least one axiom. For the convergence equation to be wrong, either the second law of thermodynamics fails, or one of five elementary properties of selection probabilities fails.

**Condition 2: A neural network cannot estimate parameters in a known equation from labeled examples**. If the convergence equation is correct, then the SFM architecture — which implements that equation by construction — reduces the learning problem to parameter estimation: fitting the embedding vectors that populate the convergence equation from training data. This is not function approximation in an unknown function class. The functional form is given; only the parameters are unknown. Parameter estimation from labeled data using gradient descent is the foundational operation of statistical learning — the basis of linear regression, logistic regression, maximum likelihood estimation, and every neural network trained since Robbins and Monro established stochastic approximation convergence guarantees (Robbins & Monro, *Annals of Mathematical Statistics*, 1951). For an SFM to fail at parameter estimation given sufficient labeled data, gradient-based optimization would have to fail on a smooth loss surface (InfoNCE) with well-characterized gradient properties.

For SFMs to fail given adequate training data, either the second law of thermodynamics must be wrong, or the foundations of statistical estimation must be broken. Both are revisable in the philosophical sense that any empirical claim is revisable. Neither is plausible given the evidence accumulated over 150 years of physics and 75 years of statistical learning theory.

### 10.2 What Remains Genuinely Uncertain

The analysis above is conditional on “sufficient training data of adequate quality.” This condition is not trivially satisfied.

Whether existing databases provide sufficient binding pair diversity for each domain is an empirical question. Antibody-antigen (SAbDab) has approximately 4,000 curated pairs; transcription factor-DNA (JASPAR + ENCODE) has orders of magnitude more; drug-target (PDBbind) has approximately 15,000. The diversity and quality of these datasets vary substantially across domains, and the convergence equation does not specify how much data is sufficient — it specifies the functional form, not the sample complexity.

Whether the cross-modal pseudo-labeling pipeline (§3.7, §9.9) can expand diversity at scale depends on the accuracy of structural co-folding models and thermodynamic calculators as pseudo-label generators. Pseudo-label noise propagates to the contrastive embedding; if noise is systematic rather than random, it can bias the learned energy landscape.

Whether the emergent bilinearity assumption (§3.1) holds quantitatively for all protein domains is untested. Bilinearity is exact for nucleic acid systems; for protein systems, it is an approximation that must be learned from data. If the approximation is poor for a given domain, the five architecture conditions are not fully satisfied, and the Level 3 alignment guarantee does not apply.

The practical questions are real. How much training data is sufficient for each domain? What retrieval accuracy is achievable at what data scale? How does performance compare to structure-first methods at matched compute budgets? How well do sequence-based predictions translate to experimental binding measurements? These are questions of degree, not of kind. The theoretical framework guarantees that the architecture implements the correct equation. Whether the available data is sufficient to populate that equation with accurate parameters is the empirical question that defines the next phase of this work.

## §11. Conclusion

The softmax attention mechanism in transformer architectures is mathematically identical to the Boltzmann distribution governing molecular binding at thermal equilibrium — the convergence equation. The Luce uniqueness theorem establishes that this is the only function satisfying the five axioms of competitive selection. Five architecture conditions — discrete contacts, bilinear energy decomposition, finite competitor pools, thermal equilibrium, and stochastic selection — bridge the convergence equation to a complete neural architecture: dual encoders, cross-attention, InfoNCE training, symmetric loss, learned temperature, and cross-attentive decoder. We term this architecture a Specificity Foundation Model and verify the five conditions across ten biological domains spanning adaptive immunity, gene regulation, RNA biology, enzymology, genome editing, pharmacology, and cell signaling.

The theoretical framework developed here extends beyond the architecture. The MRC scaling law predicts exponential convergence of retrieval accuracy with training data diversity, with a domain-specific diversity constant *D*_*C*_ determined by thermodynamic degrees of freedom at the binding interface. The affinity calibration framework derives two-parameter sufficiency for mapping contrastive scores to binding free energies, with the decoupling principle predicting that the energy landscape scales with cheap binding pair data while the unit conversion is a fixed-cost operation. The hybrid recursive learning framework prescribes cross-modal reinforcement learning with orthogonal verification — sequence-space actors verified by structure-space oracles — with a theoretical basis for collapse resistance grounded in inductive bias orthogonality. The classification of architecture-physics alignment into three levels positions SFMs as the first Level 3 case: mathematical identity between the architecture and the governing equation of the physical system.

These predictions are specific, quantitative, and falsifiable. The convergence equation and the architecture it prescribes are established by proof. Whether the predictions survive empirical testing across ten molecular recognition domains is the question that the scaling experiments, affinity calibration measurements, and cross-domain evaluations are designed to answer.

## Supplementary Note S1

### The Convergence Equation: Full Elimination Proof

#### Purpose

This note provides the complete proof that the softmax function is the unique solution to the molecular selection problem. The proof proceeds by elimination: we state five axioms of competitive selection that any selection function must satisfy, enumerate candidate function families, and show that each candidate except the exponential normalization (softmax) violates at least one property. The proof follows Luce’s Choice Axiom (Luce, *Individual Choice Behavior*, 1959) and connects to the Boltzmann distribution derivation from maximum entropy (Jaynes, *Phys. Rev*., 1957). Every step is explicit to enable verification.

### S1.1 Notation Summary

| Symbol | Definition | Domain / Constraints |
| --- | --- | --- |
| $s_j$ | Compatibility score of candidate $j$ | $s_j \in \mathbb{R}$ (unrestricted) |
| $n$ | Number of candidates in the pool | $n \geq 2$, finite |
| $P(j)$ | Selection probability of candidate $j$ | $P(j) > 0, \sum_j P(j) = 1$ |
| $\tau$ | Temperature (scaling parameter) | $\tau > 0$ |
| $g(s)$ | Positive function converting scores to weights | $g: \mathbb{R} \rightarrow \mathbb{R}^+$ |
| $E_j$ | Binding free energy of candidate $j$ | $E_j \in \mathbb{R}$ (negative = favorable) |
| $k_B T$ | Thermal energy (Boltzmann constant × temperature) | $\approx 0.6$ kcal/mol at 310 K |

### S1.2 The Selection Problem

A molecular agent (antibody, transcription factor, enzyme, guide RNA) encounters a pool of *n* candidate targets. Each candidate *j* has a compatibility score *s*_*j*_ ∈ ℝ reflecting its binding favorability. The selection function *F* maps the score vector (*s*_1_, *s*_2_, …, *s*_*n*_) to a probability distribution +*P*(1), *P*(2), …, *P*(*n*)-over the candidates. The question:

What is *F*? Is it unique?

We answer this by requiring *F* to satisfy five properties that any physically meaningful selection mechanism must possess. These properties are not hypotheses; they are constraints imposed by the physics of competitive molecular binding.

### S1.3 The Five Axiomatic Properties

#### Property 1: Positivity

For all candidates *j* and all score vectors (*s*_1_, …, *s*_*n*_): *P*(*j*) > 0

#### Physical basis

At non-zero temperature, every molecular interaction has non-zero probability. The Boltzmann distribution assigns positive probability to every energy state. A system at 310 K (physiological temperature) will sample all accessible states with non-zero frequency. A selection function that assigns zero probability to a reachable state contradicts thermodynamics.

#### What this eliminates

Any function that clips low-scoring candidates to exactly zero. This includes the argmax function (which assigns probability 1 to the highest-scoring candidate and 0 to all others) and truncated distributions that threshold below a cutoff.

#### Property 2: Normalization

For all score vectors:

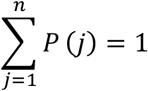

#### Physical basis

The agent binds exactly one target at a time (or, equivalently, the probability distribution over binding states sums to unity). This is a standard requirement for any probability distribution.

#### Structural consequence

Combined with Property 1, normalization forces the selection function to have the form:

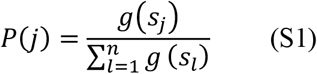

where *g*: ℝ → ℝ^+^ is a strictly positive function. The question reduces to: what is *g*?

#### Property 3: Unrestricted Domain

The function *g* must accept any real-valued input:

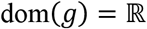

#### Physical basis

Compatibility scores are dimensionless energies *s*_*j*_ = −*E*_*j*_/*k*_B_*T*. Binding energies span the full real line: *E*_*j*_ can be positive (repulsive), zero, or negative (attractive), with no finite bound in either direction. Highly unfavorable interactions (large positive *E*_*j*_) yield large negative *s*_*j*_; highly favorable interactions (large negative *E*_*j*_) yield large positive *s*_*j*_. The function *g* must be defined for all such inputs.

#### What this eliminates

Functions with restricted domains. 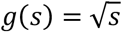 requires *s* ≥ 0. *g*(*s*) = log(*s*) requires *s* > 0. Both fail for negative compatibility scores, which occur whenever the interaction energy is repulsive.

#### Property 4: Rank Preservation (Monotonicity)

If *s*_A_> *s*_B_, then *P*(*A*) > *P*(*B*). Equivalently, *g* must be strictly increasing:

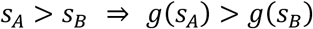

#### Physical basis

A candidate with more favorable binding energy must have higher binding probability. This is the thermodynamic principle: lower energy states are more probable at equilibrium. Any function that inverts or fails to preserve the energetic ranking contradicts the second law of thermodynamics applied to the binding equilibrium.

#### What this eliminates

Functions that are not monotonically increasing over the full real line. *g*(*s*) = *s*^2^ (or any even polynomial) fails: *g*(−3) = 9 > *g*(1) = 1, so a candidate with score −3 would have higher weight than a candidate with score +1. *g*(*s*) = sin(*s*) and all periodic functions fail: they oscillate and cannot be monotonically increasing everywhere.

#### Property 5: Independence of Irrelevant Alternatives (IIA)

For any two candidates *A* and *B* in the pool:

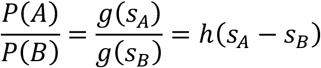

for some function *h*. That is, the probability ratio depends only on the score difference *s*_A_ – *s*_B_, not on the absolute magnitudes *s*_*A*_ and *s*_*B*_ individually, and not on any other candidates in the pool.

#### Physical basis

Adding or removing an irrelevant alternative from the pool should not change the relative odds between two existing candidates. In molecular recognition, the relative probability that an antibody binds antigen A versus antigen B depends on the difference in binding free energies ΔΔ*G* = Δ*G*_*A*_ – Δ*G*_*B*_, not on whether a third antigen C is present in the environment. This is a consequence of equilibrium thermodynamics: at equilibrium, the ratio *P*(*A*)/*P*(*B*) = exp(−(*E*_*A*_ – *E*_*B*_)/*k*_*B*_*T*) = exp(*s*_*A*_ – *s*_*B*_), which depends only on the energy difference.

#### This is the decisive property

It is the property that uniquely determines *g* to be an exponential function. The argument is as follows.

### S1.4 Derivation of the Unique Solution from IIA

We prove that *g*(*s*) = exp(*s*/*τ*) is the unique function satisfying Properties 1–5.

**Step 1 (IIA constrains the ratio)**. Properties 1–2 establish the normalized form *P*(*j*) = *g*+*s*_*j*_-/ ∑_*l*_ *g* (*s*_*l*_) with *g*: ℝ → ℝ^+^ strictly positive. Property 5 (IIA) requires that the ratio *g*(*s*_A_)/*g*(*s*_*B*_) depends only on the score difference *s*_*A*_ – *s*_*B*_. Define *ϕ*(*s*) = ln*g*(*s*). Then:

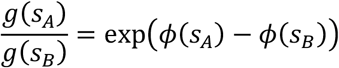

IIA requires *ϕ*(*s*_*A*_) − *ϕ*(*s*_*B*_) = *F*(*s*_*A*_ – *s*_*B*_) for some function *F*.

**Step 2 (Differentiation yields a constant)**. Differentiating *ϕ*(*s*_*A*_) − *ϕ*(*s*_*B*_) = *F*(*s*_*A*_ – *s*_*B*_) with respect to *s*_*A*_:

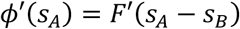

Differentiating the same identity with respect to *s*_*B*_:

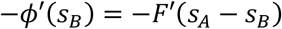

Therefore *ϕ*′(*s*_*A*_) = *ϕ*′(*s*_*B*_) for all *s*_*A*_, *s*_*B*_. The derivative *ϕ*′ is constant. Let *ϕ*′(*s*) = *c* for all *s*.

**Step 3 (Integration)**. Integrating *ϕ*′(*s*) = *c* gives *ϕ*(*s*) = *c*s + *d* for constants *c, d*. Since *g*(*s*) = exp+*ϕ*(*s*)-:

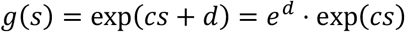

The constant *e*^*d*^ cancels in the normalization *P*(*j*) = *g*+*s*_*j*_-/ ∑_l_ *g* (*s*_l_). Property 4 (rank preservation) requires *g* strictly increasing, forcing *c* > 0. Setting *c* = 1/*τ* with *τ* > 0:

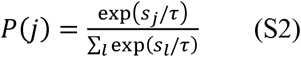

This is the softmax function with temperature *τ*. It is the unique solution.

#### Verification

Softmax satisfies all five properties:

1. exp(*s*_*j*_/*τ*) > 0 for all *s*_*j*_ ∈ ℝ, so *P*(*j*) > 0.
2. ∑_*j*_ *P* (*j*) = 1 by construction.
3. exp(*s*/*τ*) is defined for all *s* ∈ ℝ.
4. *s*_*A*_ > *s*_*B*_ implies exp(*s*_*A*_/*τ*) > exp(*s*_*B*_/*τ*), so *P*(*A*) > *P*(*B*).
5. *P*(*A*)/*P*(*B*) = exp+(*s*_*A*_− *s*_*B*_)/*τ*-, depending only on *s*_*A*_ – *s*_*B*_.

All five properties satisfied.

#### Regularity note

Step 2 assumes *ϕ* is differentiable. This can be relaxed: Aczél (1966) proved that any measurable solution to the functional equation *ϕ*(*a*) − *ϕ*(*b*) = *F*(*a* − *b*) is affine, so the conclusion *ϕ*(*s*) = *c*s + *d* holds under much weaker conditions. For physical systems where compatibility scores vary continuously with molecular configurations, differentiability is not restrictive.

### S1.5 Elimination of Candidate Function Families

The proof above establishes uniqueness constructively. For pedagogical completeness, we now verify the elimination by testing seven commonly proposed candidate families against the five properties. Each candidate is a function *g* that could be inserted into the normalization form *P*(*j*) = *g*+*s*_*j*_-/ ∑_*l*_ *g* (*s*_*l*_).

**Candidate 1: Simple (linear) normalization**

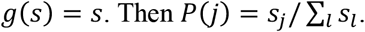

#### Fails Property 1 (Positivity)

If *s*_*j*_ < 0 (repulsive interaction), then *g*(*s*_*j*_) = *s*_*j*_ < 0, so *P*(*j*) < 0. Not a valid probability. Additionally, if ∑_*l*_*s*_*l*_ = 0, the denominator is zero and the function is undefined.

**Candidate 2: ReLU normalization**

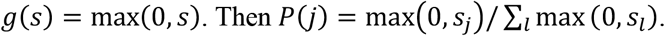

#### Fails Property 1 (Positivity)

If *s*_*j*_ ≤ 0, then *g*(*s*_*j*_) = 0, so *P*(*j*) = 0. Every candidate with non-positive compatibility score receives zero probability, violating the requirement that all candidates have non-zero binding probability at physiological temperature.

**Candidate 3: Sigmoid normalization**

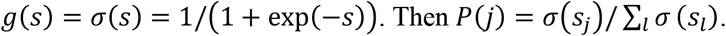

#### Satisfies Properties 1–4

*σ*(*s*) > 0 for all *s* (Property 1). Normalization is imposed by construction (Property 2). *σ* is defined for all *s* ∈ ℝ (Property 3). *σ* is strictly increasing (Property 4).

#### Fails Property 5 (IIA)

The ratio *σ*(*s*_*A*_)/*σ*(*s*_*B*_) = (1 + exp(−*s*_*B*_))/+1 + exp(−*s*_*A*_)-. Consider *s*_*A*_ − *s*_*B*_ = 1 (fixed difference). At *s*_*A*_ = 1, *s*_*B*_ = 0: ratio = (1 + 1)/(1 + *e*^−1^) = 2/1.368 = 1.462. At *s*_*A*_ = 101, *s*_*B*_ = 100: ratio = (1 + *e*^−100^)/(1 + *e*^−101^) ≈ 1.000. The ratio depends on absolute magnitudes, not only on the difference. Two candidates with the same score difference have different probability ratios depending on where they sit on the score scale. This violates the thermodynamic requirement that relative binding probabilities depend only on the free energy difference.

#### Candidate 4: Square root normalization

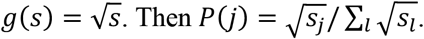

#### Fails Property 3 (Unrestricted domain)

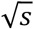 is undefined for *s* < 0. Binding energies span the full real line, so compatibility scores *s*_*j*_ = −*E*_*j*_/*k*_*B*_*T* can be arbitrarily negative. The function cannot process repulsive interactions.

#### Candidate 5: Polynomial (even power) normalization

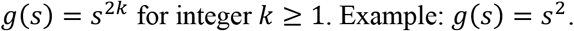

#### Fails Property 4 (Rank preservation)

*g*(−3) = 9 > *g*(1) = 1, so a candidate with score −3 receives higher weight than a candidate with score +1. The function is not monotonically increasing on ℝ. Odd-power polynomials *g*(*s*) = *s*^2*k*+1^ fail Property 1 (they take negative values for negative inputs).

#### Candidate 6: Periodic functions

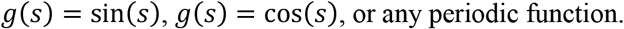

#### Fails Properties 1 and 4

Periodic functions oscillate, so they are not monotonically increasing (violates Property 4). They also take negative values (sin(π + *ϵ*) < 0), violating Property 1.

#### Candidate 7: Exponential (softmax)

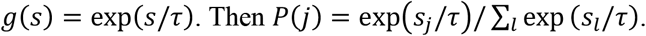

#### Satisfies all five properties

Verified in S1.4, Step 4. This is the unique survivor.

**Table S1.** Elimination of candidate function families. ✓ = satisfies property. ✗ = fails property (with explanation).

| Candidate $g(s)$ | P1 Positive | P2 Normalize | P3 Domain | P4 Monotone | P5 IIA | Status |
| --- | --- | --- | --- | --- | --- | --- |
| $s$ (linear) | × | ✓ | ✓ | ✓ | ✓ | Eliminated: $P(j) < 0$ for $s < 0$ |
| $\max(0, s)$ | × | ✓ | ✓ | × | — | Eliminated: $P(j) = 0$ for $s \leq 0$ |
| $\sigma(s)$ sigmoid | ✓ | ✓ | ✓ | ✓ | × | Eliminated: ratio depends on $\vert s\vert $ |
| $\sqrt{s}$ | $\checkmark^*$ | ✓ | × | $\checkmark^*$ | — | Eliminated: undefined for $s < 0$ |
| $s^{2k}$ | ✓ | ✓ | ✓ | × | — | Eliminated: not monotone |
| $\sin(s), \cos(s)$ | × | ✓ | ✓ | × | — | Eliminated: oscillates, goes negative |
| $\exp(s/\tau)$ | ✓ | ✓ | ✓ | ✓ | ✓ | Unique survivor |

- $\checkmark^*$ Satisfies property only on restricted domain; fails globally.

### S1.6 Elimination Table

**Table S2.** Five independent derivations of Equation S2.

| Year | Author(s) | Field | Starting point |
| --- | --- | --- | --- |
| 1877 | Boltzmann | Statistical mechanics | Maximum entropy at thermal equilibrium |
| 1948 | Shannon | Information theory | Maximum entropy distribution |
| 1957 | Jaynes | Information theory / statistical inference | Maximum entropy subject to average energy constraint |
| 1959 | Luce | Mathematical psychology | IIA axiom for rational choice behavior |
| 2017 | Vaswani et al. | Machine learning (NLP) | Scaled dot-product attention for sequence transduction |

### S1.7 Six Independent Derivations

The softmax/Boltzmann function has been derived six times across six fields with no cross-influence at the time of derivation. This convergence is itself evidence that the equation is a mathematical necessity.

Boltzmann derived the distribution from counting microstates in a gas at fixed total energy (Boltzmann, *Wien. Ber*., 1877). Shannon showed that the distribution maximizing entropy over a discrete set of outcomes subject to a mean-value constraint takes the same exponential form (Shannon, *Bell Syst. Tech. J*., 1948). Jaynes unified these two results, re-deriving the Boltzmann distribution as the least-biased probability assignment consistent with a known average energy (Jaynes, *Phys. Rev*., 1957). Luce derived the same functional form from choice axioms, without reference to physics — his starting point was the IIA property of human decision-making (Luce, *Individual Choice Behavior*, 1959). Vaswani et al. introduced scaled dot-product attention for neural machine translation, selecting softmax as the attention function (Vaswani et al., *NeurIPS*, 2017). The mathematical form is identical in all five cases. The Boltzmann distribution has been applied to molecular binding equilibria since the development of statistical thermodynamics, and is the standard formalism for transcription factor occupancy (Bintu *et al., Curr. Opin. Genet. Dev*., 2005) and nucleic acid duplex stability (SantaLucia, *PNAS*, 1998).

The contribution of the present work is not a sixth derivation but the identification that these five derivations converge on the same equation, the proof via Luce’s uniqueness theorem that no other equation is possible (S1.4), and the demonstration that this identity — combined with five architecture conditions (Supplementary Note S2) — prescribes a complete neural architecture for molecular recognition across ten biological domains (Reddy, *bioRxiv*, 2026).

### S1.8 Connection to the Boltzmann Distribution

The selection problem (S1.2) and the Boltzmann distribution describe the same mathematics with different vocabulary. We make the correspondence explicit.

In statistical mechanics, a system at thermal equilibrium occupies microstates with probabilities proportional to exp(−*E*/*k*_*B*_*T*), where *E* is the microstate energy. The partition function *Z* = ∑_*j*_ exp +−*E*_*j*_/*k*_*B*_*T*-normalizes the distribution:

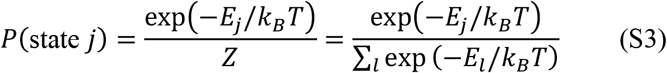

Define the compatibility score *s*_*j*_ = −*E*_*j*_/*k*_*B*_*T*. Substituting into Equation S3:

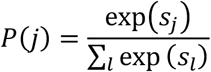

This is Equation S2 with *τ* = 1. The temperature *τ* in the general softmax corresponds to *k*_*B*_*T* in the physical system. Low temperature (*τ* → 0) concentrates probability on the highest-scoring candidate (strongest binder); high temperature (*τ* → ∞) gives a uniform distribution (all candidates equally likely).

**The Boltzmann distribution is the softmax function applied to negative energies scaled by temperature**.

This is not an analogy. It is an algebraic identity: 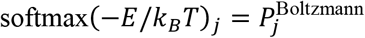. The uniqueness proof (S1.4) shows that this is the only function consistent with the physics of competitive molecular selection.

### S1.9 Connection to Transformer Attention

Transformer attention computes:

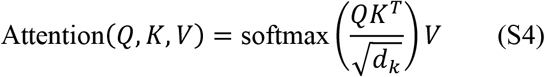

where *Q, K, V* are query, key, and value matrices, and 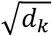 is a scaling factor (Vaswani et al., *NeurIPS*, 2017). The correspondence to molecular recognition is described in Table 3.

**Table S3.** Correspondence between molecular selection and transformer attention.

| Molecular recognition | Transformer attention | Mathematical role |
| --- | --- | --- |
| Agent features (paratope, DBD, seed) | Query vector $q$ | “What am I looking for?” |
| Target features (epitope, motif, site) | Key vector $k$ | “What do I offer?” |
| Binding energy $E = -q \cdot k$ | Dot-product score $q \cdot k$ | Compatibility |
| Thermal energy $k_B T$ | Scale factor $\sqrt{d_k}$ | Selection sharpness (temperature) |
| Binding probability $P(j \vert i)$ | Attention weight $A_{ij}$ | Selection probability |
| Information at binding interface | Value vector $v$ | What is transferred upon selection |

The identity is exact when binding energy decomposes as a dot product (§3.1 Condition 2, main text). This holds exactly for position weight matrix systems (TF–DNA, miRNA–mRNA, CRISPR gRNA– DNA) and in learned embedding space for systems with non-modular binding energy (Ab–Ag, TCR– pMHC, enzyme–substrate). See Supplementary Note S2 for the full treatment of this distinction.

### S1.10 Scope and Limitations

#### What the proof establishes

The softmax function is the unique solution to the competitive molecular selection problem under five physically motivated axioms. The proof is mathematical, not empirical. It holds for any system satisfying the five properties, regardless of the specific molecules involved.

#### Assumption: differentiability

The derivation in S1.4, Step 2 assumes *ϕ* = ln(*g*) is differentiable. This can be relaxed: Aczél showed that any measurable solution to Cauchy’s functional equation is linear (Aczél, *Lectures on Functional Equations and Their Applications*, 1966), so the conclusion *ϕ*(*s*) = *cs* + *d* holds under much weaker regularity conditions. For physical systems where scores vary continuously with molecular configurations, differentiability is not a restrictive assumption.

#### The IIA assumption

IIA (Property 5) is the most restrictive axiom and the one most likely to be violated in non-equilibrium settings. If the system is not at thermal equilibrium — for example, if binding is kinetically trapped — then the relative odds between two candidates may depend on other candidates (through kinetic competition for shared intermediates). The present framework applies to equilibrium binding. Extension to non-equilibrium systems (e.g., Phase 2 of CRISPR R-loop propagation, which involves irreversible conformational changes) would require relaxing IIA.

#### Temperature is the only free parameter

The five properties determine the functional form completely. The only freedom is the temperature *τ*, which sets the sharpness of selection. In physical systems, *τ* = *k*_*B*_*T* is measured. In learned embedding spaces (CALM, TCR-SFM, eSFM), *τ* is a hyperparameter. The proof does not determine *τ*; it determines everything else.

### S1.11 Prior Art and Novelty

The mathematical content of this note — the uniqueness of the exponential normalization function under Properties 1–5 — is a classical result that has been discovered independently in at least six fields. We make the attribution explicit.

#### Mathematical psychology

Luce (1959) proved that the Independence of Irrelevant Alternatives axiom (Property 5), combined with positivity and normalization, uniquely determines the exponential choice rule. This is Luce’s Choice Axiom. It is the foundational result of mathematical choice theory and has been a textbook theorem for over sixty years (Luce, *Individual Choice Behavior*, 1959).

#### Econometrics

McFadden (1974) applied Luce’s result to derive the multinomial logit model for discrete economic choices — the probability that a consumer selects option *j* from a choice set is *P*(*j*) = exp(*V*_*j*_)/ ∑_*l*_ exp (*V*_*l*_), where *V*_*j*_ is the utility of option *j*. This is Equation S2 with utility in place of compatibility score. McFadden received the 2000 Nobel Prize in Economics partly for this work (McFadden, *Econometrica*, 1974). The multinomial logit model is taught in every economics PhD program.

#### Statistical mechanics

Boltzmann (1877) and Jaynes (1957) derived the same function from maximum entropy at thermal equilibrium. This derivation does not use the IIA axiom explicitly; instead, it maximizes the Shannon entropy (1948) of the probability distribution subject to a constraint on the average energy. The result is the same: *P*(*j*) = exp(−*E*_*j*_/*k*_*B*_*T*)/*Z*. The Boltzmann distribution is undergraduate physics.

#### Functional equations

The step in our derivation (S1.4, Step 2) where IIA forces *ϕ*′(*s*) = constant is an instance of Cauchy’s functional equation. Aczél (1966) proved that any measurable solution to Cauchy’s equation is linear, establishing the result under weaker regularity conditions than differentiability (Aczél, *Lectures on Functional Equations*, 1966). This is a classical result in functional analysis.

#### What is known vs. what is new

The mathematical theorem (Properties 1–5 ⇒ softmax) is known. What is new is the interpretive chain from this theorem to neural architecture design for molecular recognition. Specifically:

i. No prior work has stated the five properties as axiomatic constraints on *molecular binding* and derived softmax as the unique selection function for molecular recognition. Biophysicists use the Boltzmann distribution as an established result but do not frame it as a uniqueness theorem from choice axioms. The IIA property, which is decisive (it is what eliminates sigmoid and all other candidates), has not previously been articulated as a physical requirement of competitive molecular binding.
ii. No prior work has drawn the consequence that the uniqueness of softmax *determines the neural architecture* of a foundation model. Luce did not design neural networks. McFadden did not train protein language models. The insight that a sixty-year-old uniqueness theorem in mathematical psychology implies that the transformer attention mechanism is the physics-optimal architecture for predicting molecular binding specificity — and that any other architecture trains against the convergence equation — is the contribution of this paper and the previous work on computational convergence (Reddy, *bioRxiv*, 2026).
iii. The elimination table (Table S1) and the explicit candidate-by-candidate verification (S1.5) are pedagogical, not mathematically novel. Their purpose is to make the uniqueness proof verifiable by a reader (or automated system) that has not encountered Luce’s theorem. The goal is verification, not originality.

We state this clearly because the strength of the argument rests on the fact that the mathematics is established, not speculative. If the uniqueness of softmax were a new conjecture, the downstream claims about architecture derivation would be provisional. Because the uniqueness is a proven theorem with independent derivations in four fields, the downstream claims are deductive consequences of known mathematics applied to a new domain.

### S1.12 Summary of Proof

**Table S4.** Summary: from five properties to the convergence equation.

| Step | Starting point | Result |
| --- | --- | --- |
| 1 | Positivity + Normalization | $P(j) = g(s_j) / \sum g(s_i)$, with $g: \mathbb{R} \rightarrow \mathbb{R}^+$ |
| 2 | Unrestricted domain | Eliminates $\sqrt{s}$, $\log(s)$, and all functions with restricted domains |
| 3 | Rank preservation | Eliminates even polynomials, periodic functions; $g$ must be strictly increasing |
| 4 | IIA | Forces $g(s)/g(s') = h(s - s')$, which requires $g(s) = \exp(s/\tau)$ |
| 5 | Verification | $\exp(s/\tau)$ satisfies all five properties. Unique survivor. |
| 6 | Change of variables | $s_j = -E_j / k_B T$ recovers Boltzmann. $s_j = q \cdot k_j$ recovers attention. |

### S1.13 Formal Statement

#### Theorem (Luce’s Choice Axiom, restated for molecular selection)

*Let P*: ℝ^*n*^ → *Δ*^*n*−1^ be a function mapping score vectors to probability distributions over *n* ≥ 2 candidates. Suppose *P satisfies: (1) P*(*j*) > 0 *for all* j and all score vectors (positivity); (2) ∑_*j*_ *P* (*j*) = 1 *(normalization); (3) P* is defined for all *s*_*j*_ ∈ ℝ (unrestricted domain); (4) *s*_*A*_ > *s*_*B*_ ⇒ *P*(*A*) *> P(B)* (rank preservation); and (5) *P*(*A*)/*P*(*B*) *depends only on s*_*A*_ – *s*_*B*_ (IIA). Then there exists *τ* > 0 such that:

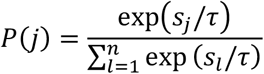

**and this is the unique such function**.

### S1.14 Verification Checklist

□ Five axiomatic properties are physically motivated and explicitly stated
□ Property 5 (IIA) is the decisive property that determines the functional form
□ Derivation in S1.4 uses IIA to establish Cauchy’s equation, yielding φ(s) = cs + d
□ Exponentiation recovers g(s) = exp(s/τ), with normalization constant canceling
□ All seven candidate families are tested; only exp(s/τ) survives all five properties
□ Sigmoid satisfies P1–P4 but fails P5 (IIA) — the key elimination
□ Six independent derivations (Boltzmann, Jaynes, Luce, Vaswani) yield same equation
□ Boltzmann–softmax identity is exact: softmax(−E/k_*B*_T) = P(Boltzmann)
□ Transformer attention correspondence is exact when energy is bilinear
□ Temperature τ is the only free parameter; functional form is fully determined
□ IIA limitation acknowledged: proof applies to equilibrium systems only

## Supplementary Note S2

### Structural Conditions for Architecture-Physics Alignment

#### Purpose

This note provides the detailed justification for each of the five structural conditions introduced in §3 of the main text, verified across all ten molecular recognition domains. For each condition, we state the physical requirement, explain why it is necessary for the Boltzmann-to-Softmax attention identity, and document how each domain satisfies it. We give particular attention to the distinction between given and emergent bilinearity (Condition 2), which has architectural and scaling consequences, and to domain-specific masking and bias mechanisms — chromatin accessibility masking (tSFM), PAM-site masking (crisprSFM), and concentration-weighted attention bias (eSFM) — that arise naturally from the structural conditions. Every claim is explicit to enable verification.

### S2.1 Overview: From Convergence Equation to Architecture

Supplementary Note S1 proved that the softmax function is the unique selection mechanism for competitive molecular binding (Eq. 1, main text). This is unconditional — it holds for any system at thermal equilibrium selecting among finite alternatives. The structural conditions in this note are additional requirements that enable the further identification of softmax selection with transformer attention. The distinction is:

#### Softmax alone

(Eq. 1) says: selection probabilities are exponentially weighted by compatibility scores. This requires only equilibrium and finite alternatives.

#### Softmax attention

(Eq. 3) says: the compatibility scores are dot products of learned feature vectors, and the output is a weighted sum of value vectors. This requires all five structural conditions.

A system satisfying only the first two conditions could still use softmax selection but would need a non-attention architecture to compute the scores. A system satisfying all five admits a full dual-encoder transformer architecture — what we call a Specificity Foundation Model.

### S2.2 Condition 1: Discrete Intermolecular Contacts

#### Statement

Binding occurs through a finite set of contacts between specific positions on two separate molecules.

Why it is necessary. Transformer attention operates on sequences of discrete tokens. Each position in the query attends to each position in the key. For the attention mechanism to correspond to molecular binding, the binding interaction must likewise decompose into position-specific contacts between two distinct molecular entities. Continuous-field interactions (e.g., long-range electrostatic gradients between charged surfaces) do not decompose into discrete contact pairs and cannot be represented as position-to-position attention weights.

#### Verification across ten domains

Ab–Ag (CALM): Binding occurs through 10–30 residue–residue contact pairs between CDR loops (paratope) and epitope surface (Chothia & Lesk, *Journal of Molecular Biology*, 1987). Contact is defined by distance threshold (≤5 Å between heavy atoms). The contacts are discrete (each residue either contacts or does not contact a partner residue) and positionally defined (CDR1, CDR2, CDR3 loops on VH and VL chains).

TCR–pMHC (TCR-SFM): Structurally homologous to Ab–Ag. TCR CDR loops (α and β chains) contact peptide residues in the MHC groove through 15–25 contact pairs (Rudolph et al., *Annual Review of Immunology*, 2006). The CDR3 loops dominate peptide contact; CDR1/CDR2 primarily contact MHC helices.

TF–DNA (tSFM): Transcription factor DNA-binding domains (DBDs) contact specific nucleotide positions within a binding motif (typically 6–20 bp). Each position-nucleotide pair (4 possibilities per position) constitutes a discrete contact. The number of contacts equals the motif length *L* times 4 possible nucleotides — a finite, well-enumerated set (Stormo, *Annual Review of Biophysics*, 2000).

miRNA–mRNA (miR-SFM): MicroRNA targeting operates through Watson–Crick base-pairing between the ∼22 nt miRNA guide and complementary sites in the mRNA 3’UTR. Each nucleotide position in the miRNA pairs (or mispairs) with a specific nucleotide in the target. The seed region (positions 2–8) dominates target recognition (Bartel, *Cell*, 2009). Contacts are discrete and positionally defined.

Enzyme–substrate (eSFM): Enzyme–substrate binding occurs through contacts between active-site residues and specific atoms or functional groups on the substrate. The Mechanism and Catalytic Site Atlas (M-CSA; Ribeiro et al., *Nucleic Acids Res*., 2018) enumerates catalytic residues for characterized enzymes. Contact pairs are discrete: each catalytic residue interacts with a specific substrate moiety.

CRISPR gRNA–DNA (crisprSFM): The ∼20 nt guide RNA spacer hybridizes with the DNA protospacer through positional base-pairing. Each of 20 positions constitutes a discrete contact. Mismatch tolerance varies with position: PAM-proximal (seed, positions 1–8) mismatches are highly disruptive; PAM-distal mismatches are increasingly tolerated (Hsu et al., *Nature Biotechnology*, 2013). Contacts are discrete and have a defined positional gradient.

### S2.3 Condition 2: Bilinear Energy Decomposition

#### Statement

The binding energy between agent *A* and target *j* decomposes as a dot product of feature vectors:

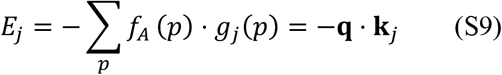

where **q** encodes the agent’s positional preferences and **k**_*j*_ encodes the target’s positional properties.

#### Why it is necessary

The dot product **q** ⋅ **k** is the compatibility score in transformer attention. Substituting a dot-product energy into the Boltzmann distribution yields: *P*(*j*) = exp(**q** ⋅ **k**_*j*_/*τ*)/ ∑_*l*_ exp (**q** ⋅ **k**_*l*_/*τ*), which is the scaled dot-product attention equation (Vaswani et al., *NeurIPS*, 2017). Without bilinear energy decomposition, the Boltzmann selection probability still holds (S1), but the connection to *dot-product attention specifically* does not. Condition 2 is what connects the convergence equation to the transformer architecture.

#### Given vs. emergent bilinearity

This is the most important distinction across the ten domains. It has consequences for temperature mode, data efficiency, and scaling behavior.

**Given bilinearity** means the dot-product decomposition exists in the raw physical data, not in a learned representation. The binding energy is literally a sum of independent positional contributions, each expressible as a product of an agent feature and a target feature. The agent and target feature vectors exist in interpretable physical coordinates.

#### TF–DNA (given)

The position weight matrix (PWM) formalism expresses TF–DNA binding energy as:

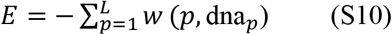

where *w*(*p, b*) is the weight for nucleotide *b* at position *p*, and *L* is the motif length. This is a dot product: the TF preference vector (a 4*L*-dimensional vector encoding *w*(*p, b*) for all positions and nucleotides) dotted with the one-hot-encoded DNA sequence (a 4*L*-dimensional vector with ones at the observed nucleotides). The PWM IS the dot product in raw sequence space (Stormo, *Annual Review of Biophysics*, 2000). The bilinearity is given by the physics, not learned by the model.

#### miRNA–mRNA (given)

Under the nearest-neighbor thermodynamic model, RNA duplex binding energy decomposes as a sum of per-position contributions (with stacking corrections between adjacent positions): *ΔG* ≈ ∑_*p*_ *Δ G* (bp_*p*_) (SantaLucia, *PNAS*, 1998). The base-pairing energy at each position is a product of the miRNA nucleotide identity and the target nucleotide identity — a bilinear form. Stacking corrections (S2.8) represent deviations from pure bilinearity that map to relative positional encoding.

#### CRISPR gRNA–DNA (given)

Binding energy decomposes as a position-weighted sum: *ΔG* = ∑_*p*_ *w* (*p*) ⋅ *ΔG* (bp_*p*_), where *w*(*p*) is a position-dependent weight reflecting the seed-to-distal mismatch tolerance gradient (Hsu et al., *Nature Biotechnology*, 2013). This is a weighted dot product between the guide RNA’s positional preferences and the protospacer’s base-pair properties. The weighting is biophysically determined (not learned), making the bilinearity given.

**Emergent bilinearity** means the raw binding energy does not decompose as a positionally independent sum. Shape complementarity, cooperative interactions, and long-range electrostatics contribute to binding in ways that are not bilinear in raw coordinates. However, neural network encoders can learn a feature representation in which the effective energy does decompose as a dot product. The bilinearity emerges in the learned embedding space.

#### Ab–Ag (emergent)

Antibody–antigen binding involves shape complementarity, electrostatics, hydrophobic packing, and hydrogen bonding across a large, geometrically complex interface. These do not decompose into positionally independent contributions in raw coordinates. CALM learns an embedding space in which the effective binding score sim(**q, k**) approximates the true binding affinity as a dot product (Lee et al., *bioRxiv*, 2026).

#### TCR–pMHC (emergent)

Same as Ab–Ag. TCR–pMHC binding involves shape complementarity and non-additive interactions across the CDR–peptide–MHC interface. Bilinearity is emergent in embedding space.

#### Enzyme–substrate (emergent)

Enzyme–substrate binding involves shape complementarity, electrostatic steering, and transition-state stabilization at the active site. These interactions are geometrically complex and non-additive in raw coordinates. Bilinearity is emergent in embedding space.

#### Drug–target (emergent)

Small molecule binding to protein targets involves van der Waals contacts, hydrogen bonds, electrostatic interactions, and desolvation penalties across a geometrically defined binding pocket. The energy does not decompose as a positionally independent sum in raw coordinates. Bilinearity is emergent in the learned embedding space of atom-level molecular fingerprints.

#### Peptide–MHC (emergent)

Peptide binding to MHC grooves involves anchor residue contacts at defined pockets (P2, P9 for class I) and distributed contacts along the groove. While anchor positions contribute semi-independently, the full binding energy includes cooperative effects between anchor and non-anchor positions. Bilinearity is emergent.

#### Receptor–ligand (emergent)

Receptor-ligand binding at cell surface interfaces involves protein-protein or protein-small molecule contacts with shape complementarity and electrostatic steering. Bilinearity is emergent in embedding space, analogous to Ab–Ag.

#### RNA-binding protein–RNA (emergent, partially given for RRM domains)

Some RNA-binding proteins (particularly those with RNA recognition motifs, RRMs) recognize short sequence elements with partial positional independence, suggesting partially given bilinearity analogous to PWMs. Other RBP classes (KH domains, zinc fingers) recognize more complex structural elements where bilinearity is fully emergent.

**Table S7.**
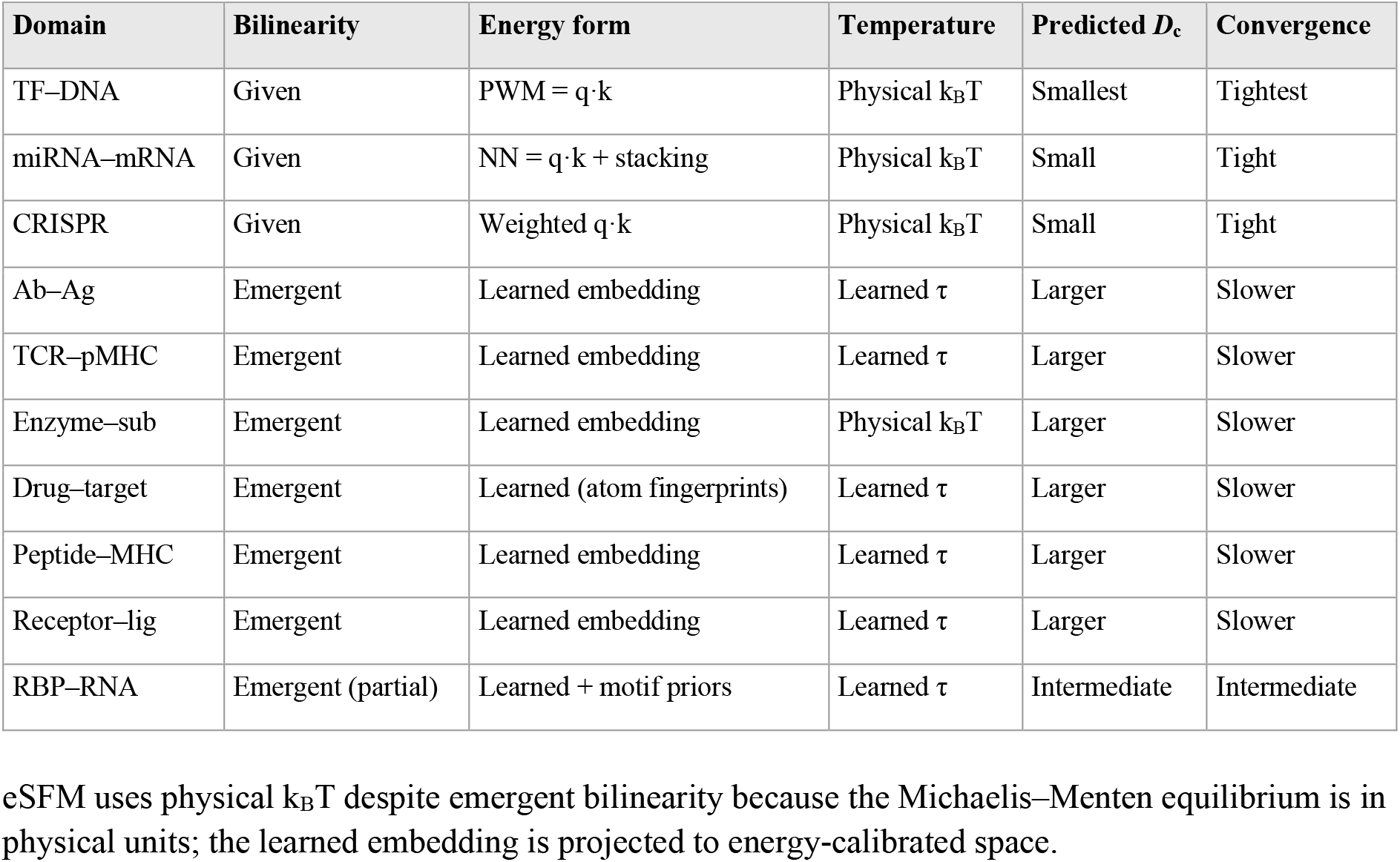
Given vs. emergent bilinearity across ten domains.

| Domain | Bilinearity | Energy form | Temperature | Predicted $D_c$ | Convergence |
| --- | --- | --- | --- | --- | --- |
| TF–DNA | Given | PWM = $\mathbf{q} \cdot \mathbf{k}$ | Physical $k_B T$ | Smallest | Tightest |
| miRNA–mRNA | Given | NN = $\mathbf{q} \cdot \mathbf{k}$ + stacking | Physical $k_B T$ | Small | Tight |
| CRISPR | Given | Weighted $\mathbf{q} \cdot \mathbf{k}$ | Physical $k_B T$ | Small | Tight |
| Ab–Ag | Emergent | Learned embedding | Learned $\tau$ | Larger | Slower |
| TCR–pMHC | Emergent | Learned embedding | Learned $\tau$ | Larger | Slower |
| Enzyme–sub | Emergent | Learned embedding | Physical $k_B T$ | Larger | Slower |
| Drug–target | Emergent | Learned (atom fingerprints) | Learned $\tau$ | Larger | Slower |
| Peptide–MHC | Emergent | Learned embedding | Learned $\tau$ | Larger | Slower |
| Receptor–lig | Emergent | Learned embedding | Learned $\tau$ | Larger | Slower |
| RBP–RNA | Emergent (partial) | Learned + motif priors | Learned $\tau$ | Intermediate | Intermediate |

eSFM uses physical k_*B*_T despite emergent bilinearity because the Michaelis–Menten equilibrium is in physical units; the learned embedding is projected to energy-calibrated space.

#### S2.3.1 Scaling prediction from bilinearity type

Table S7 records a prediction: systems with given **bilinearity** will have smaller scaling constants *D*_1_ and faster convergence than systems with emergent **bilinearity**. We now derive this prediction from classical estimation theory.

##### The learning problem differs by bilinearity type

In a system with given bilinearity (TF–DNA, miRNA–mRNA, CRISPR), the functional form of the energy is known: *E* = −**q** ⋅ **k** in raw physical coordinates. The model does not need to discover that binding energy is a dot product. It needs only to learn the parameter values that populate the known equation — the weights of the PWM, the stacking energies, the positional tolerance gradient. This is **parameter estimation** within a known model class.

In a system with emergent bilinearity (Ab–Ag, TCR–pMHC, enzyme–substrate), the raw binding energy does not decompose as a dot product. The model must learn a nonlinear embedding *ϕ*:sequence → ℝ^*d*^ such that sim(*ϕ*(agent), *ϕ*(target)) -G / k_*B*_T. It must discover both the functional form (the embedding that makes energy bilinear) and the parameter values within that form. This is **function approximation** — a strictly harder problem.

##### Estimation theory predicts the data requirement

The classical result from statistical estimation theory is that the sample complexity of parameter estimation scales with the number of free parameters *p*, while the sample complexity of nonparametric function approximation scales with the intrinsic dimensionality of the function class (Wasserman, *All of Nonparametric Statistics*, 2006). For a model with *p* parameters estimated within a known functional form, the mean squared error decreases as:

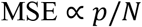

where *N* is the number of training samples. For nonparametric estimation (unknown functional form), the rate is:

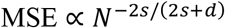

where *s* is the smoothness of the function and *d* is its intrinsic dimensionality. The parametric rate (1/*N*) is always faster than the nonparametric rate (*N*^−*α*^ with *α* < 1). More unknowns require more data. This is a fundamental result, not specific to molecular recognition.

##### Application to SFMs

In the exponential scaling law *R* = *R*_max_ × (1 − exp(−*D*/*D*_*c*_)) (Eq. 6, main text), the scaling constant *D*_1_ reflects the effective number of unknowns the model must learn from training data. For given-bilinearity systems, the unknowns are the physical parameters of the energy function (PWM weights, stacking energies). For emergent-bilinearity systems, the unknowns include both the embedding function and the parameters within it. Therefore:

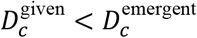

The prediction is quantitative. If *D*_*c*_ correlates with thermodynamic degrees of freedom at the binding interface (Section 4.3, main text), then given-bilinearity systems should cluster at lower *D*_*c*_ values and emergent-bilinearity systems at higher *D*_*c*_ values on the *D*_*c*_ vs. degrees-of-freedom plot (Figure 8, main text). The separation between the two clusters is a direct measurement of the data cost of learning the embedding function.

##### Falsifiable test

If a given-bilinearity system (e.g., tSFM) shows *D*_*c*_ comparable to or larger than an emergent-bilinearity system (e.g., CALM), one of two conclusions follows: either the bilinearity classification is wrong (the system’s energy does not in fact decompose as a dot product in raw coordinates), or additional complexity not captured by the bilinearity condition (e.g., severe epistasis, conformational flexibility) dominates the learning problem. Both are informative. The prediction provides a diagnostic, not just a forecast.

##### The matched filter analogy

The distinction between given and emergent bilinearity parallels the distinction between a matched filter and an adaptive filter in signal processing. A matched filter (North, *RCA Technical Report*, 1943) knows the signal waveform and estimates only its amplitude and phase — parameter estimation. An adaptive filter must learn the waveform from data — function approximation. The matched filter requires orders of magnitude less data for the same detection performance. Given-bilinearity SFMs are matched filters for molecular recognition; emergent-bilinearity SFMs are adaptive filters. Both converge to the same solution; the matched filter gets there faster.

### S2.4 Condition 3: Finite Competitor Pools

#### Statement

The agent selects among a finite number of candidate targets.

#### Why it is necessary

The softmax function operates on a finite sum: 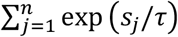. This is a discrete sum, not an integral. If the competitor pool were continuous (infinitely many targets at infinitesimally different energies), the sum would become an integral and the mathematical form would change. Finite pools ensure the softmax denominator is well-defined and computable.

#### Verification

Ab–Ag: antigens in serum or tissue environment (finite, though large). TCR–pMHC: peptides presented by MHC molecules on cell surfaces (finite, ∼10^4^–10^5^ distinct pMHC per cell). TF– DNA: accessible genomic binding sites (∼10^4^–10^6^ per cell type, determined by chromatin state). miRNA–mRNA: target sites in the 3’UTR transcriptome (∼10^4^ per miRNA seed family). Enzyme– substrate: metabolites in the cellular environment (finite, ∼10^3^–10^4^ distinct small molecules). CRISPR gRNA–DNA: PAM-adjacent genomic sites (∼10^6^–10^8^ per genome, depending on PAM frequency).

All pools are finite.

### S2.5 Condition 4: Thermal Equilibrium

#### Statement

The system operates at a well-defined temperature, and binding occupancy follows the Boltzmann distribution.

#### Why it is necessary

The Boltzmann distribution is the equilibrium distribution. If the system is not at equilibrium — if binding is kinetically trapped, or if the system has not had time to reach steady state — the occupancy probabilities deviate from Boltzmann. The softmax identity requires equilibrium.

#### Verification across ten domains

#### Ab–Ag: Assumed

Standard assumption for antibody–antigen binding kinetics. Binding and unbinding rates (*k*_on_, *k*_off_) typically reach equilibrium on timescales of minutes to hours, well within the timescale of immune responses.

#### TCR–pMHC: Assumed

TCR–pMHC interactions are transient (*K*_*D*_ in the micromolar range, *t*_1/2_ of seconds to minutes) and rapidly reach equilibrium. Serial triggering models assume repeated equilibrium binding events (Rudolph et al., *Annual Review of Immunology*, 2006).

#### TF–DNA: Validated

TF occupancy at genomic sites follows Boltzmann statistics, as demonstrated experimentally and theoretically (Bintu et al., *Current Opinion in Genetics & Development*, 2005). The thermodynamic models of gene regulation are quantitatively successful, confirming equilibrium.

#### miRNA–mRNA: Validated

miRNA–mRNA duplex formation reaches thermodynamic equilibrium, as evidenced by the quantitative success of nearest-neighbor thermodynamic models in predicting duplex stability (SantaLucia, *PNAS*, 1998; Bartel, *Cell*, 2009).

#### Enzyme–substrate: Validated (Michaelis–Menten)

The Michaelis–Menten model assumes rapid equilibrium between enzyme and substrate: *E* + *S* ⇌ *ES* → *E* + *P* (Michaelis & Menten, *Biochem. Z*., 1913). The equilibrium assumption is standard and experimentally validated for the binding step. Catalysis (product formation) is irreversible, but the selection step (which substrate binds the active site) is at equilibrium.

#### CRISPR gRNA–DNA: Validated (Phase 1)

CRISPR target recognition proceeds in two phases. Phase 1: PAM interrogation and seed hybridization — reversible, rapid, at equilibrium. Phase 2: R-loop propagation past ∼12 matched positions — irreversible, committing to cleavage (Sternberg et al., *Nature*, 2014). The SFM models Phase 1 selection probability only. The equilibrium boundary is well-defined.

#### Drug–target: Validated (*K*_*d*_)

Small molecule binding to protein targets reaches thermodynamic equilibrium on timescales of seconds to minutes. Dissociation constants (*K*_*d*_ values) measured by isothermal titration calorimetry, surface plasmon resonance, and fluorescence polarization are equilibrium measurements. The SFM framework applies to equilibrium binding; kinetic selectivity (residence time effects) is not modeled.

#### Peptide–MHC: Assumed

Peptide loading onto MHC molecules is mediated by chaperones (tapasin for class I, HLA-DM for class II) and involves kinetic editing. The final peptide-MHC complex, once formed, is at thermodynamic equilibrium with a stability measurable by thermal denaturation assays. The SFM models the equilibrium complex, not the loading pathway.

#### Receptor–ligand: Validated (*K*_*d*_)

Receptor-ligand interactions at the cell surface reach binding equilibrium, with *K*_*d*_ values spanning picomolar to micromolar ranges depending on the receptor-ligand pair. Equilibrium binding constants are routinely measured by radioligand binding assays and surface plasmon resonance.

#### RNA-binding protein–RNA: Validated

RBP-RNA interactions reach equilibrium binding, with *K*_*d*_ values measurable by electrophoretic mobility shift assay (EMSA), fluorescence anisotropy, and CLIP-seq enrichment quantification.

### S2.6 Condition 5: Stochastic Selection

#### Statement

Binding is probabilistic at physiological temperature, not deterministic.

#### Why it is necessary

If selection were deterministic (the agent always binds the single best target), the selection function would be argmax, not softmax. The Boltzmann distribution assigns non-zero probability to all accessible states at *T* > 0 (Property 1, S1). At physiological temperature (310 K, *k*_*B*_*T* ≈ 0.6 kcal/mol), thermal fluctuations are comparable to many binding energy differences, ensuring stochastic selection. The softmax function with finite temperature captures this stochasticity; argmax (*τ* → 0) does not.

#### Verification

All ten systems operate in aqueous solution at ∼310 K. Thermal energy (*k*_*B*_*T* ≈ 0.6 kcal/mol) is comparable to typical binding energy differences between cognate and non-cognate pairs (1–10 kcal/mol for most molecular recognition systems). This ensures that selection is probabilistic, not deterministic. Even the strongest binder does not have probability 1; even weak binders have non-zero probability. This is a universal feature of molecular recognition at physiological temperature.

### S2.7 Domain-Specific Masking Mechanisms

Three domains require attention masks that restrict the candidate pool before softmax selection is applied. These masks do not violate the structural conditions; they enforce them. A masked position is physically inaccessible and therefore not a valid candidate. Masking ensures that the softmax denominator sums only over accessible alternatives.

#### S2.7.1 Chromatin accessibility masking (tSFM)

In eukaryotic cells, genomic DNA is packaged into chromatin. Nucleosome-wrapped regions are physically inaccessible to transcription factors. Only open chromatin regions — identified by ATAC-seq or DNase-seq — are available for TF binding. The chromatin accessibility mask is:

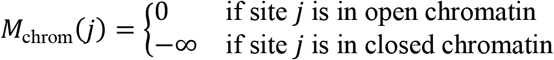

The mask is added to the attention logits before softmax: *P*(*j*) = softmax (**q** ⋅ **k**_*j*_/*τ* + *M*_chrom_(*j*)). Sites with *M* = −∞ receive zero probability after softmax, exactly as in transformer attention masking. The mask is **cell-type-specific**: different cell types have different open chromatin landscapes (ENCODE Project Consortium, *Nature*, 2012). The same TF may have different effective target pools in different cell types, implemented by different masks.

This is a structural correspondence, not a metaphor. Transformer attention masking prevents the model from attending to certain positions; chromatin masking prevents the TF from binding certain genomic sites. The mathematical implementation is identical.

#### S2.7.2 PAM-site masking (crisprSFM)

Cas9 (and other Cas proteins) cannot interrogate a genomic site unless a valid protospacer adjacent motif (PAM) is present immediately adjacent to the protospacer (Jinek et al., *Science*, 2012). PAM recognition is a prerequisite that gates all subsequent steps of target interaction. A site without a valid PAM has zero probability of binding, regardless of spacer–protospacer complementarity. The PAM mask is:

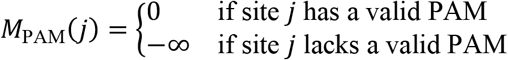

Multi-PAM support: SpCas9 recognizes NGG, SaCas9 recognizes NNGRRT, Cas12a recognizes TTTV. The mask function parameterizes the PAM requirement per Cas variant: *M*_PAM_(*j*; Cas) = 0 if the PAM motif for the specified Cas protein is present at site *j*, −∞ otherwise. Unlike chromatin masking, PAM masking is **sequence-determined and cell-type-invariant**: the PAM is a DNA sequence feature, not an epigenetic one.

#### S2.7.3 Concentration-weighted attention bias (eSFM)

In vivo, enzyme–substrate selection depends not only on binding affinity (*ΔG*) but also on substrate concentration. At unequal concentrations, the selection probability is:

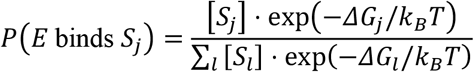

Taking the logarithm of the numerator: ln ([*S*_*j*_] ⋅ exp(−*ΔG*_*j*_/*k*_*B*_*T*)) = ln[*S*_*j*_] − *ΔG*_*j*_/*k*_*B*_*T*. Therefore, the effective attention logit is:

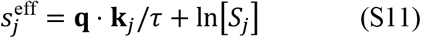

The ln[*S*_*j*_] term is an additive bias to the attention logits, implementable as a bias vector from metabolomics data or pathway models. This is not an approximation; it is the exact Boltzmann probability when concentrations are unequal. The bias is optional: when all substrate concentrations are equal ([*S*_*j*_] = [*S*_*l*_] for all *j, l*), the ln[*S*_*j*_] terms cancel in the softmax and the standard form (Eq. 3) is recovered.

### S2.8 Deviations from Pure Bilinearity: Epistatic Corrections

Three given-bilinearity systems (TF–DNA, miRNA–mRNA, CRISPR gRNA–DNA) exhibit known deviations from pure positional independence: the binding energy at position *p* depends not only on the identity of the nucleotide at position *p* but also on the identity of the nucleotide at adjacent positions. These are epistatic (dinucleotide or nearest-neighbor stacking) effects. We show that these deviations do not break the convergence identity. They refine it: each class of epistatic correction maps to a specific, known attention architecture variant.

#### S2.8.1 The pure bilinear baseline

Under pure positional independence, the binding energy is:

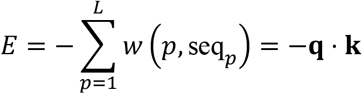

This is a dot product (Eq. S9). The softmax selection probability is vanilla dot-product attention. Each position contributes independently to the total energy.

#### S2.8.2 Dinucleotide corrections in TF–DNA binding

Real TF–DNA binding includes dinucleotide effects: the energy contribution of the base at position *p* depends on the identity of the base at position *p* + 1 (and vice versa). The corrected energy is:

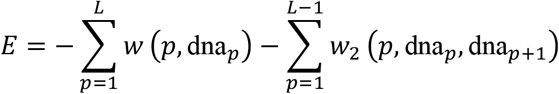

The first sum is the PWM (dot product). The second sum is the dinucleotide correction — a function of two adjacent positions.

The dinucleotide correction *w*_2_+*p*, dna_*p*_, dna_*p*+1_) depends on the identity of the nucleotide at position *p* and the identity of the nucleotide at position *p* + 1. This is a pairwise, position-dependent energy term. In transformer architecture, a position-dependent pairwise bias added to the attention logits is the definition of relative positional encoding (Shaw et al., *NAACL*, 2018). The correspondence is:

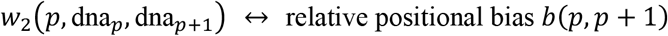

The dinucleotide energy matrix (4 × 4 at each adjacent position pair) maps directly to the relative positional encoding matrix in the DNA encoder. The correction does not change the selection mechanism (still softmax); it changes the score computation within the encoder.

#### S2.8.3 Nearest-neighbor stacking in miRNA–mRNA and CRISPR gRNA–DNA

RNA duplex stability follows the nearest-neighbor thermodynamic model (SantaLucia, *PNAS*, 1998). The binding free energy is:

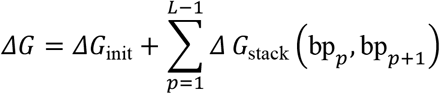

where *ΔG*_stack_ (bp_*p*_, bp_*p*+1_) is the stacking free energy of adjacent base-pair doublets. This is structurally identical to the TF–DNA dinucleotide correction: a pairwise, position-dependent energy term depending on the identity of the nucleotides at positions *p* and *p* + 1. The same correspondence applies: nearest-neighbor stacking energies map to relative positional encoding in the RNA or DNA encoder.

For CRISPR gRNA–DNA, the relevant thermodynamic parameters are those for DNA:RNA hybrid duplexes, which differ from RNA:RNA parameters (Sugimoto et al., *Biochemistry*, 1995). The functional form is identical; only the numerical values of *ΔG*_stack_ change. The architectural correspondence (stacking → relative PE) is unchanged.

#### S2.8.4 Why epistatic corrections do not break the convergence

Two observations establish this.

**First**, the Boltzmann distribution (Eq. 1) holds regardless of the energy functional form. Whether *E* is a pure dot product, includes dinucleotide corrections, or has arbitrary higher-order terms, the selection probability is:

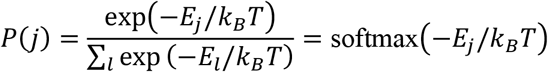

The selection mechanism is always softmax. The five-property uniqueness proof (S1) does not depend on the form of the energy. It depends only on the properties of the selection function. Epistatic corrections modify the compatibility score *s*_*j*_; they do not modify the selection function.

**Second**, the corrections map to known, well-characterized attention variants. The mapping is summarized in Table S9.

**Table S9.**
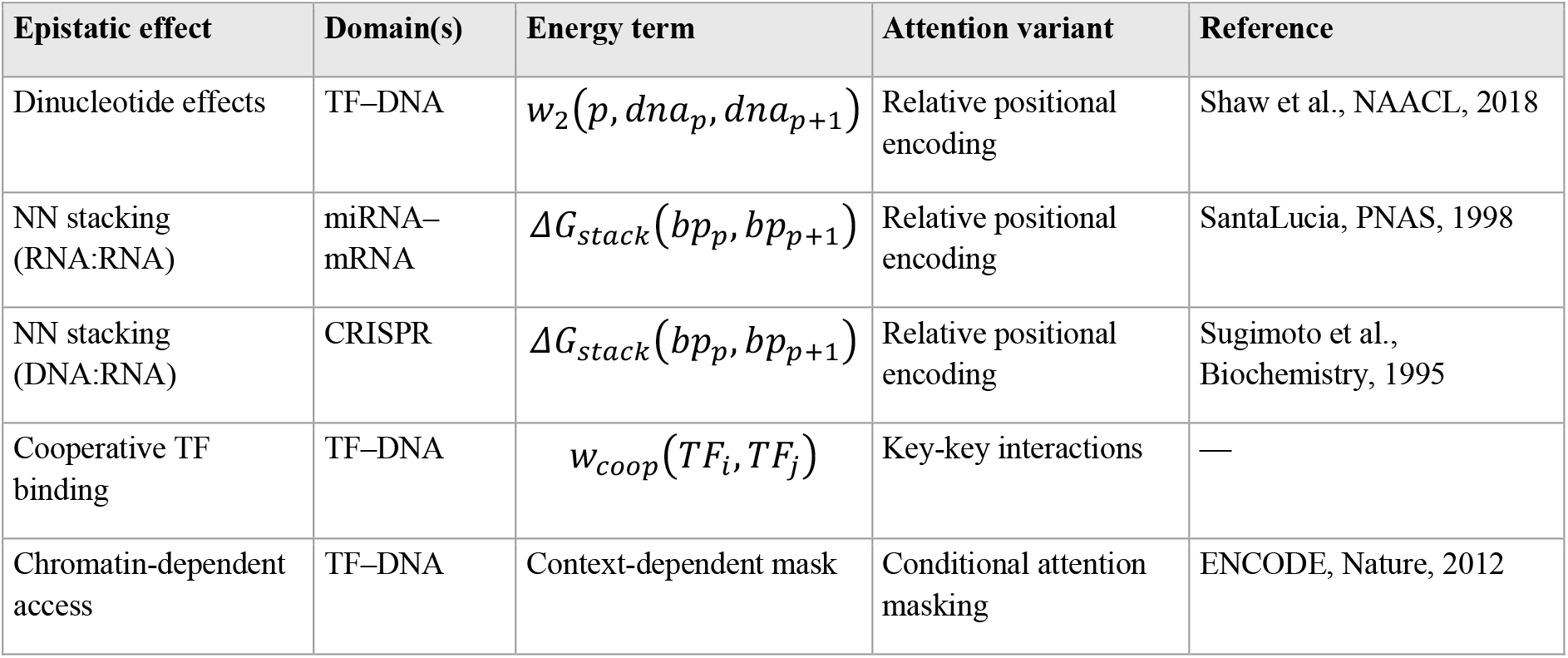
Epistatic corrections and their attention architecture correspondences.

| Epistatic effect | Domain(s) | Energy term | Attention variant | Reference |
| --- | --- | --- | --- | --- |
| Dinucleotide effects | TF–DNA | $w_2(p, dna_p, dna_{p+1})$ | Relative positional encoding | Shaw et al., NAACL, 2018 |
| NN stacking (RNA:RNA) | miRNA–mRNA | $\Delta G_{\text{stack}}(bp_p, bp_{p+1})$ | Relative positional encoding | SantaLucia, PNAS, 1998 |
| NN stacking (DNA:RNA) | CRISPR | $\Delta G_{\text{stack}}(bp_p, bp_{p+1})$ | Relative positional encoding | Sugimoto et al., *Biochemistry*, 1995 |
| Cooperative TF binding | TF–DNA | $w_{\text{coop}}(TF_i, TF_j)$ | Key-key interactions | — |
| Chromatin-dependent access | TF–DNA | Context-dependent mask | Conditional attention masking | ENCODE, Nature, 2012 |

The first three rows are pairwise positional effects — all map to relative positional encoding. Cooperative TF binding (two TFs binding adjacent sites with mutual stabilization) maps to key-key interactions in attention. Chromatin-dependent accessibility (treated in S2.7.1) is a conditional mask.

#### S2.8.5 Architectural implication

The epistatic corrections are not noise, failure, or limitation. They are architectural discoveries. Each correction identifies which specific attention variant is physics-optimal for the domain. A pure PWM system (no epistasis) uses vanilla dot-product attention. A system with dinucleotide effects uses dot-product attention with relative positional encoding. A system with cooperative binding uses attention with key-key interactions. Biology specifies the attention variant; the convergence identity specifies the selection mechanism.

This has a practical consequence for SFM design: the choice of positional encoding in the DNA/RNA encoder is not a hyperparameter to be tuned — it is determined by the biophysics of the domain. Systems with significant nearest-neighbor stacking effects (miRNA–mRNA, CRISPR gRNA–DNA) should use relative positional encoding; systems without (or with weak epistasis) need not.

### S2.9 Summary Table

**Table S8.** Complete structural conditions verification. All five conditions, all ten domains.

| Condition | Ab–Ag | TCR–pMHC | TF–DNA | miR–mRNA | Enz–sub | CRISPR | Drug–tgt | Pep–MHC | Rec–lig | RBP–RNA |
| --- | --- | --- | --- | --- | --- | --- | --- | --- | --- | --- |
| 1. Discrete | Res–res | Res–res | Res–nt | Nt–nt | Res–atom | Nt–nt | Atom–res | Res–res | Res–res | Res–nt |
| 2. Bilinear | Emergent | Emergent | Given (PWM) | Given (NN) | Emergent | Given (wt.) | Emergent | Emergent | Emergent | Emergent |
| 3. Finite | Serum Ags | Peptides | Genomic | 3'UTR | Metabol. | PAM sites | Proteome | Processed | Ligands | Transcripts |
| 4. Equilib. | Assumed | Assumed | Validated | Validated | Val. (M–M) | Val. (Ph.1) | Val. (K<sub>d</sub>) | Assumed | Val. (K<sub>d</sub>) | Validated |
| 5. Stochastic | ✓ | ✓ | ✓ | ✓ | ✓ | ✓ | ✓ | ✓ | ✓ | ✓ |
| Temperature | Learned $\tau$ | Learned $\tau$ | Phys. k<sub>B</sub>T | Phys. k<sub>B</sub>T | Phys. k<sub>B</sub>T | Phys. k<sub>B</sub>T | Learned $\tau$ | Learned $\tau$ | Learned $\tau$ | Learned $\tau$ |
| Masking | None | None | Chromatin | None | Conc. bias | PAM | None | None | None | None |

### S2.10 Verification Checklist

□ All ten domains satisfy Condition 1 (discrete contacts): contact type specified per domain
□ Given bilinearity verified for TF–DNA (PWM), miRNA–mRNA (NN model), CRISPR (weighted dot product)
□ Emergent bilinearity justified for Ab–Ag, TCR–pMHC, enzyme–substrate, drug–target, peptide– MHC, receptor–ligand, RBP–RNA (learned embedding)
□ Given bilinearity predicts tighter convergence (smaller *D*_*c*_) than emergent bilinearity
□ All ten domains have finite competitor pools; pool sizes estimated per domain
□ Equilibrium validated or assumed per domain; CRISPR Phase 1/Phase 2 boundary explicit
□ Stochastic selection justified by physiological temperature (k_*B*_T ≈ 0.6 kcal/mol at 310 K)
□ Chromatin masking for tSFM: cell-type-specific, ATAC-seq/DNase-seq data, M = −∞ for closed
□ PAM masking for crisprSFM: sequence-determined, cell-type-invariant, multi-PAM support
□ Concentration bias for eSFM: ln[S_*j*_] additive to attention logits, exact from Boltzmann
□ Epistatic corrections do not break convergence; they specify attention variants
□ Temperature mode follows from bilinearity type: given → physical, emergent → learned

## Supplementary Note S3

### Derivation of Contrastive Training Objective from Molecular Selection

#### Purpose

This note provides the complete mathematical derivation of the InfoNCE contrastive learning objective (Oord et al., *arXiv*, 2018) from the physics of competitive molecular selection. We show that InfoNCE is the negative log-likelihood of the Boltzmann selection probability derived in §2 of the main text and Supplementary Note S1. The derivation establishes that InfoNCE is not one of many possible training objectives — it is the unique objective that trains a model to reproduce the convergence equation of molecular recognition. We then extend the derivation to the symmetric (bidirectional) case used in all ten SFMs, derive the role of hard negatives, and establish the correspondence to clonal selection in adaptive immunity. Every step is explicit to enable verification.

### S3.1 Notation Summary

| Symbol | Definition | Domain |
| --- | --- | --- |
| $\mathbf{q}$ | Agent embedding (query) | $\mathbf{q} \in \mathbb{R}^d$ |
| $\mathbf{k}_f$ | Correct target embedding (positive key) | $\mathbf{k}_f \in \mathbb{R}^d$ |
| $\mathbf{k}_{c_i}$ | Non-cognate competitor embedding (negative key i) | $\mathbf{k}_{c_i} \in \mathbb{R}^d$ |
| $M$ | Number of competitors (negatives) | $M \geq 1$ |
| $\text{sim}(\cdot, \cdot)$ | Similarity function (cosine or dot product) | $\text{sim}: \mathbb{R}^d \times \mathbb{R}^d \rightarrow \mathbb{R}$ |
| $\tau$ | Temperature (selection stringency) | $\tau > 0$ |
| $P(f\vert \mathbf{q})$ | Selection probability for correct target | $P \in (0,1)$ |
| $\mathcal{L}$ | Training loss | $\mathcal{L} \geq 0$ |
| $N$ | Mini-batch size | $N \geq 2$ |

### S3.2 The Selection Scenario

A molecular agent encounters a pool of candidates. Exactly one candidate is the correct (cognate) target; the remaining *M* candidates are non-cognate competitors. This scenario arises in every molecular recognition domain:

In antibody–antigen recognition, a B cell receptor encounters multiple antigens; the immune system must select cells that bind the pathogenic antigen over self or irrelevant antigens. In transcription factor–DNA binding, a TF selects its cognate motif from among all accessible genomic sites. In CRISPR, a guide RNA–Cas9 complex selects the on-target protospacer from among all PAM-adjacent sites in the genome. In each case, the physics is identical: competitive selection among finite alternatives at thermal equilibrium.

The question: What training objective causes a model to learn the parameters that correctly reproduce this selection process?

### S3.3 The Selection Probability

From §2 of the main text (Eq. 1) and Supplementary Note S1, the unique selection probability satisfying the five axiomatic properties is the softmax (Boltzmann) function. Applied to the agent– target selection scenario:

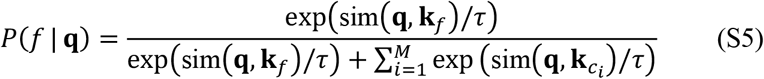

where sim(**q, k**) is the compatibility score between agent and target in the shared embedding space. This is the probability that a correctly parameterized model assigns to the cognate target. It is not a modeling assumption — it follows from the convergence equation.

### S3.4 From Selection Probability to Training Loss

#### Step 1: Maximum likelihood

Given a training set of *N*_train_ agent–target pairs 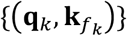, the model parameters should maximize the joint probability of correctly selecting all cognate targets:

where *θ* denotes the encoder parameters. This is standard maximum likelihood estimation.

#### Step 2: Negative log-likelihood

Taking the logarithm (which preserves the optimum) and negating (to convert maximization to minimization):

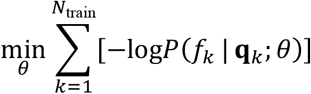

#### Step 3: Substituting the selection probability

Inserting Equation S5 into the negative log-likelihood for a single training pair:

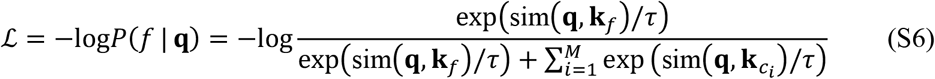

#### Step 4: Expand

Using −log(*a*/*b*) = −log*a* + log*b*:

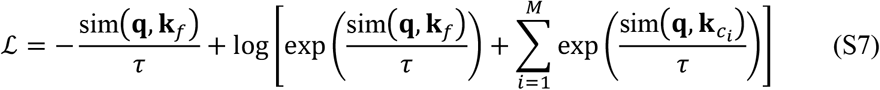

The first term rewards high similarity to the correct target. The second term (log-sum-exp) penalizes high similarity to any candidate, including the correct one. Together they enforce discrimination: the model must score the correct target higher than all competitors.

#### Step 5: Identification

Equation S6 is the InfoNCE loss (Oord et al., *arXiv*, 2018). The derivation is complete:

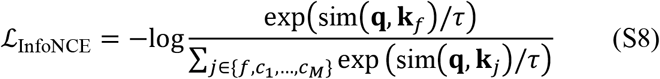

InfoNCE is the negative log-likelihood of the Boltzmann selection probability. No alternative derivation was used; no design choice was made. The loss follows from the convergence equation by maximum likelihood.

### S3.5 Why InfoNCE and Not Other Contrastive Losses

Several alternative contrastive objectives exist in the machine learning literature. The derivation above shows why InfoNCE is physics-optimal for molecular recognition and why alternatives are suboptimal.

#### Triplet loss

The triplet loss (Schroff et al., *CVPR*, 2015) enforces a margin between positive and negative similarities:

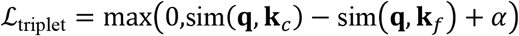

This considers only one negative at a time and enforces a fixed margin *α*. It does not model the full competitive pool. The Boltzmann distribution assigns probability based on the relative scores of all competitors simultaneously (Eq. S5). A loss function that considers only pairwise comparisons discards information about the full competitor landscape. Triplet loss is not a maximum likelihood estimator for the Boltzmann selection probability.

#### Binary cross-entropy

Binary cross-entropy treats each pair independently as a binary classification (binding vs. non-binding):

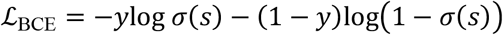

where *σ*(*s*) is the sigmoid function. Supplementary Note S1 proved that sigmoid violates the Independence of Irrelevant Alternatives (Property 5). Binary cross-entropy models absolute binding probability, not relative selection among competitors. It cannot capture the competitive nature of molecular recognition, where an agent’s binding to one target depends on the available alternatives.

#### Margin-based contrastive loss

The margin-based loss (Hadsell et al., *CVPR*, 2006) uses a fixed distance threshold rather than softmax competition:

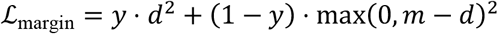

This imposes a hard geometric boundary (margin *m*) rather than the soft probabilistic competition of the Boltzmann distribution. It does not include temperature and does not compute selection probabilities. The physics requires soft competition (Property 1: all candidates have non-zero probability); a hard margin contradicts this.

#### Summary

InfoNCE is the unique loss function that is the negative log-likelihood of the Boltzmann selection probability over a finite competitor pool. All other contrastive losses either (a) ignore the full competitor pool (triplet), (b) use a function that violates the axiomatic properties (BCE/sigmoid), or (c) impose hard geometric boundaries inconsistent with thermodynamic selection (margin). Training with InfoNCE trains the model to reproduce the convergence equation. Training with any other objective trains the model to reproduce a different, non-physical target.

### S3.6 The Symmetric (Bidirectional) Loss

Binding affinity is a property of the agent–target complex, not of either partner individually. The free energy *ΔG* of an antibody–antigen complex is the same whether computed from the antibody’s perspective (which antigen does this antibody select?) or the antigen’s perspective (which antibody does this antigen select?). This physical symmetry motivates the symmetric loss used in CLIP (Radford et al., *ICML*, 2021) and in all ten SFMs.

For a mini-batch of *N* agent–target pairs, define the *N* × *N* similarity matrix *S*_*ij*_ = sim(**q**_*i*_, **k**_*j*_)/*τ*. The diagonal entries are correct pairs; off-diagonal entries are in-batch negatives. The symmetric loss is:

where:

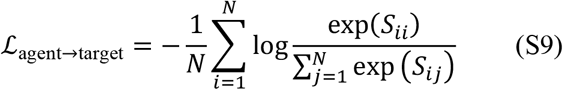

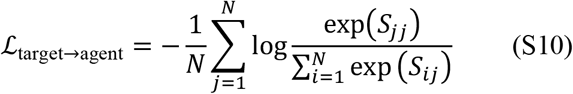

The first term asks: for each agent, can the model identify the correct target from the batch? The second term asks: for each target, can the model identify the correct agent? Both are InfoNCE applied along different axes of the similarity matrix. Lee et al. (*bioRxiv*, 2026) confirmed that CALM achieves symmetric retrieval performance (overlapping confidence intervals between Ag→Ab and Ab→Ag directions), validating that the symmetric loss produces a balanced embedding space.

### S3.7 In-Batch Negatives as the Competitor Pool

In the SFM training procedure, the competitor pool is the mini-batch itself: for each agent–target pair, all other targets in the batch serve as negatives. With batch size *N*, each training example has *M* = *N* − 1 competitors.

This connects to the Boltzmann distribution in a precise way. In equilibrium thermodynamics, the denominator of the Boltzmann distribution (the partition function *Z*) sums over all accessible states. In training, the partition function is approximated by the batch:

Larger batches provide a better approximation to the true partition function (the sum over all possible targets). This is why batch size matters for contrastive learning: it controls the quality of the competitor pool. In molecular recognition, this corresponds to the diversity of the local molecular environment. An antibody in serum encounters many distinct antigens simultaneously; a transcription factor in the nucleus encounters many competing DNA sites. The effective batch size of the biological system is the number of simultaneous competitors.

### S3.8 Hard Negatives and AIRE

Not all negatives are equally informative. A non-cognate target that is easily distinguished from the correct target (high score difference) contributes little gradient. A non-cognate target that is difficult to distinguish (similar score to the correct target) contributes the dominant gradient signal. These are **hard negatives**.

In the Boltzmann framework, the gradient of InfoNCE with respect to the similarity score for negative *c*_*i*_ is proportional to *P*(*c*_*i*_|**q**) — the selection probability of that negative. Hard negatives have high *P*(*c*_*i*_|**q**) and therefore contribute the most to learning. Easy negatives have *P*(*c*_*i*_|**q**) ≈ 0 and contribute approximately zero gradient.

In adaptive immunity, this principle is implemented by the AIRE protein (autoimmune regulator), which drives ectopic expression of tissue-restricted self-antigens in thymic medullary epithelial cells (Anderson et al., *Science*, 2002). AIRE presents the immune system’s hardest negatives: self-antigens that resemble foreign pathogens closely enough to be dangerous if not explicitly filtered. The computational parallel is exact: AIRE provides hard negatives for thymic selection; hard negative mining provides hard negatives for contrastive training. Both ensure that the model (receptor or SFM) learns to discriminate at the boundaries of the specificity landscape.

Each SFM domain has domain-specific hard negative strategies, described in Section 3 of the main text. These are not ad hoc — they follow from the same principle: provide competitors that are maximally confusable with the correct target to maximize gradient information per training step.

### S3.9 The Temperature Parameter

The temperature *τ* in the InfoNCE loss (Eq. S8) has the same mathematical role as physical temperature *k*_*B*_*T* in the Boltzmann distribution. Its effect on training is derived from thermodynamics.

#### Low *τ*. (stringent selection)

The selection probability concentrates on the highest-scoring candidate. Small differences in similarity produce large differences in probability. The model is rewarded for making fine discriminations. In molecular terms: low temperature sharpens binding specificity; the system strongly favors the best binder.

#### High *τ*. (permissive selection)

The selection probability spreads across many candidates. Even modest differences in similarity produce small differences in probability. The model is rewarded for maintaining a balanced embedding space. In molecular terms: high temperature permits promiscuous binding; many targets are nearly equally probable.

In SFMs with given bilinearity (tSFM, miR-SFM, crisprSFM), *τ* can be fixed at the physical value *k*_*B*_*T* ≈ 0.6 kcal/mol at 310 K, because the compatibility scores are in physical energy units. In SFMs with emergent bilinearity (CALM, TCR-SFM, eSFM), *τ* is learned because the compatibility scores are in arbitrary embedding units. CALM initializes *τ* = 0.2 (Lee et al., *bioRxiv*, 2026).

### S3.10 Correspondence to Clonal Selection

The derivation above is general: it applies to any molecular recognition system satisfying the five structural conditions (Section 2, main text). For the two adaptive immune SFMs (CALM and TCR-SFM), there is an additional biological correspondence. The InfoNCE loss is mathematically identical to the selection probability governing clonal selection in the adaptive immune system (Burnet, *BMJ*, 1957; Reddy, *bioRxiv*, 2026).

**Table S5.** Correspondence between immune selection and InfoNCE training.

| Immune selection | InfoNCE training | Mathematical role |
| --- | --- | --- |
| Receptor (BCR or TCR) | Anchor embedding $q$ | Agent |
| Foreign antigen (pathogen) | Positive embedding $k_f$ | Correct target |
| Self-antigens | Negative embeddings $k_c$ | Competitors |
| Binding affinity $a(r, p)$ | Similarity $\text{sim}(q, k)$ | Compatibility score |
| Selection probability $P(\text{select } f)$ | Softmax probability | Eq. S5 = Eq. 1 |
| Clonal deletion (death of wrong) | Gradient descent (minimize loss) | Optimization |
| AIRE (tissue-restricted self-antigens) | Hard negative mining | Informative competitors |
| Selection stringency | Temperature $\tau$ | Distribution sharpness |
| Germinal center competitor pool | Mini-batch | Effective denominator |

This correspondence was established in the computational convergence paper (Reddy, *bioRxiv*, 2026). The present note demonstrates that it extends beyond adaptive immunity: the InfoNCE objective is physics-optimal for all ten molecular recognition domains, including those with no evolutionary connection to the immune system (TF–DNA, enzyme–substrate, CRISPR). The immune system is the proof of concept; the convergence equation is general.

### S3.11 Summary of Derivation

**Table S6.** Summary: from the convergence equation to the training objective.

| Step | Starting point | Result |
| --- | --- | --- |
| 1 | Boltzmann selection (Eq. 1) | $P(f \mid q) = \text{softmax}(\text{sim}(q, k_f) / \tau)$ over competitor pool |
| 2 | Maximum likelihood | Maximize $\prod P(f_k \mid q_k)$ over training set |
| 3 | Negative log | $\mathcal{L} = -\log P(f \mid q) = \text{InfoNCE}$ (Eq. S8) |
| 4 | Physical symmetry of binding | Symmetric loss: average of agent→target and target→agent |
| 5 | Gradient analysis of InfoNCE | Hard negatives dominate learning; AIRE is the biological implementation |
| 6 | Boltzmann temperature | $\tau$ controls selection stringency; physical $k_B T$ or learned |

#### Final result

The convergence equation (Eq. 1) determines both the architecture (softmax attention) and the training objective (InfoNCE). These are not independent design choices. They are the selection mechanism and its likelihood function, respectively. A model trained with InfoNCE over Boltzmann-weighted embeddings is trained to reproduce the physics of molecular recognition.

### S3.12 Verification Checklist

□ Selection probability (Eq. S5) follows from the convergence equation (Eq. 1)
□ InfoNCE (Eq. S8) is the negative log of the selection probability — pure algebra
□ Maximum likelihood estimation is the standard, not a design choice
□ Triplet loss is suboptimal: considers only one negative, not the full pool
□ Binary cross-entropy is suboptimal: uses sigmoid, which violates IIA (S1)
□ Margin-based loss is suboptimal: imposes hard boundary, violates positivity (S1)
□ Symmetric loss follows from physical symmetry of binding free energy
□ In-batch negatives approximate the partition function (Boltzmann denominator)
□ Hard negatives have highest gradient contribution: ∂ℒ/∂s_{c_i} ∝ P(c_i | q)
□ AIRE correspondence is biologically accurate (Anderson et al., Science, 2002)
□ Temperature τ has identical role in Boltzmann distribution and InfoNCE
□ Correspondence table entries are accurate for immune and non-immune domains

## Supplementary Note S4

### Affinity Calibration: From Contrastive Scores to Binding Free Energies

#### Purpose

This note provides the complete mathematical derivation connecting SFM contrastive scores to experimentally measured binding affinities. We derive the two-parameter affinity calibration framework (*ΔG* = *ws* + *b*) from the convergence equation, show that each parameter has a precise physical interpretation, and establish the decoupling principle that separates the cheap scaling axis (binding pairs) from the expensive calibration axis (measured affinities). We ground the derivation in the biophysics of how binding affinities are measured experimentally, because the calibration connects the learned embedding space directly to these measurements.

### S4.1 The Binding Equilibrium

When a molecular agent (antibody, enzyme, receptor) and its target (antigen, substrate, ligand) are mixed in solution, they form reversible complexes:

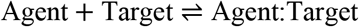

Association occurs at rate *k*_on_ (units: M^−1^s^−1^) and dissociation at rate *k*_off_ (units: s^−1^). At equilibrium, the rates balance, defining the equilibrium dissociation constant:

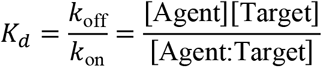

*K*_*d*_ has units of molar concentration (M). A smaller *K*_*d*_ indicates tighter binding: less free agent and target remain at equilibrium. Typical ranges span picomolar (10^-12^ M) for high-affinity therapeutic antibodies to micromolar (10^-6^ M) for weak or transient interactions.

### S4.2 From *K*_*d*_ to *ΔG*: The Boltzmann Connection

The equilibrium constant *K*_*d*_ is related to the binding free energy by a fundamental thermodynamic relationship:

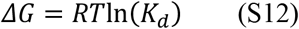

where *R* = 1.987 × 10^−3^ kcal/(mol·K) is the gas constant and *T* is temperature in Kelvin. This relationship is exact at equilibrium — it is a direct consequence of the Boltzmann distribution, not an approximation.

#### Where the Boltzmann distribution enters

At thermal equilibrium, the probability that a system occupies a state with energy *E* is proportional to exp(−*E*/*k*_*B*_*T*). For the binding reaction, there are two states: bound (free energy *ΔG* below the unbound reference) and unbound (energy = 0 by convention). The ratio of Boltzmann probabilities is:

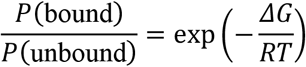

This ratio equals the association constant *K*_*a*_ = 1/*K*_*d*_ = exp(−*ΔG*/*RT*), which gives Equation S12 by rearrangement. Every *K*_*d*_ measurement — by any technique — is a measurement of the Boltzmann probability ratio between bound and unbound states.

#### Numerical scale

At *T* = 310 K (physiological temperature), *RT* = 0.616 kcal/mol. A strong antibody-antigen interaction with *K*_*d*_ = 1 nM has *ΔG* = 0.616 × ln(10^−9^) = −12.7 kcal/mol. A weak interaction with *K*_*d*_ = 1 µM has *ΔG* = 0.616 × ln(10^−6^) = −8.5 kcal/mol. The entire dynamic range of biologically relevant binding — from weak to strong — spans approximately −6 to −15 kcal/mol, a range of about 15 *RT* units.

### S4.3 How Binding Affinities Are Measured

Three experimental techniques provide the *K*_*d*_ values that serve as calibration targets.

#### Surface plasmon resonance (SPR)

One binding partner is immobilized on a gold-coated sensor chip; the other flows over the surface in solution. Binding increases the mass on the chip, detected as a shift in the angle of reflected polarized light. The instrument records a sensorgram — response units (RU) versus time — from which *k*_on_, *k*_off_, and *K*_*d*_ = *k*_off_/*k*_on_ are extracted. SPR is the gold-standard measurement for protein-protein binding kinetics and is the source of most *K*_*d*_ values in the Structural Antibody Database (SAbDab; Dunbar *et al., Nucleic Acids Res*., 2014). The equilibrium response at multiple analyte concentrations follows the Langmuir isotherm:

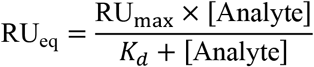

which is a direct consequence of Boltzmann equilibrium — the same equation as the Michaelis-Menten isotherm for enzyme kinetics (Michaelis & Menten, *Biochem. Z*., 1913).

#### Isothermal titration calorimetry (ITC)

ITC measures the heat released or absorbed upon binding as one partner is titrated into the other. The technique directly measures *ΔG, ΔH* (enthalpy), and *TΔS* (entropy) of binding — the complete thermodynamic signature — without requiring immobilization or labeling. ITC provides the most thermodynamically rigorous *K*_*d*_ measurements but requires large quantities of purified material.

#### Bio-layer interferometry (BLI)

BLI measures binding kinetics by detecting changes in the interference pattern of light reflected from a fiber-optic biosensor tip. One partner is immobilized on the tip; the other is in solution. BLI provides *k*_on_, *k*_off_, and *K*_*d*_ with higher throughput than SPR, though with somewhat lower precision.

All three techniques measure the same physical quantity: the equilibrium constant *K*_*d*_, which is the Boltzmann probability ratio between bound and unbound states (Equation S12).

### S4.4 The Contrastive Score as Binding Free Energy

In the convergence equation (§2 of the main text; Supplementary Note S1), the selection probability of target *j* by agent *i* is:

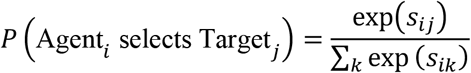

where *s*_*ij*_ = sim(**q**_*i*_, **k**_*j*_)/*τ* is the contrastive similarity score — the dot product of the agent and target embeddings, divided by the learned temperature.

The Boltzmann distribution gives the same probability in physical units:

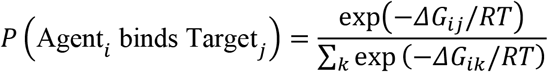

These are the same equation if and only if:

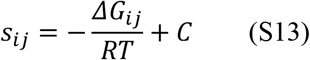

where *C* is a constant that cancels in the softmax normalization. The contrastive score IS the binding free energy in units of *RT*, up to an additive constant corresponding to the thermodynamic reference state.

This is not a modeling assumption — it is a deductive consequence of the convergence equation. If the SFM architecture implements the convergence equation (Level 3 alignment, §4 of the main text), and if the convergence equation is the Boltzmann distribution (proven in Supplementary Note S1), then the learned contrastive scores must be proportional to binding free energies. The constant *C* drops out in retrieval (ranking is preserved regardless of additive constants), which is why SFMs achieve good retrieval without ever seeing a *K*_*d*_ value. But recovering absolute *ΔG* requires calibrating out *C*.

### S4.5 Two-Parameter Sufficiency

Rearranging Equation S13:

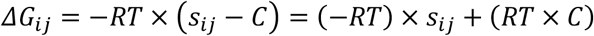

Defining *w* = −*RT* (or, more generally, a learned scaling factor absorbing the embedding-to-energy unit conversion) and *b* = *RT* × *C* (the reference-state energy):

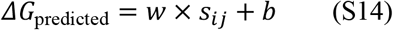

This is a two-parameter linear regression. Two parameters suffice because the convergence equation prescribes a linear relationship between contrastive scores and binding free energies. The regression is not “teaching” the SFM to predict affinity — the SFM has already learned the energy landscape through contrastive training. The regression performs a unit conversion: from embedding-space units (arbitrary, determined by encoder architecture and training dynamics) to physical units (kcal/mol, determined by the thermodynamic reference frame).

#### Sufficiency test

If two parameters are truly sufficient, adding higher-order terms should not improve the fit. Fitting both a linear model (*ΔG* = *ws* + *b*, 2 parameters) and a quadratic model (*ΔG* = *as*^2^ + *ws* + *b*, 3 parameters) and comparing by BIC (Kass & Raftery, *J. Am. Stat. Assoc*., 1995) provides a direct test. A *Δ*BIC > 10 favoring the linear model would constitute strong evidence for two-parameter sufficiency.

### S4.6 Physical Interpretation of the Regression Parameters

#### The slope *w* is the temperature / unit conversion factor

In the Boltzmann identity, *s*_*ij*_ = −*ΔG*_*ij*_/*RT*, so *ΔG*_*ij*_ = −*RT* × *s*_*ij*_. If the SFM embedding space were perfectly calibrated in energy units, *w* = −*RT* = −0.616 kcal/mol at 310 K. In practice, the embedding-space dot product has an arbitrary scale determined by the projection head dimension, L2 normalization, and the learned temperature *τ*. The slope *w* absorbs this scaling. Its sign should be negative (higher similarity = more negative *ΔG* = tighter binding), and its magnitude encodes how many kcal/mol correspond to one embedding-space unit of similarity.

#### The intercept *b* is the thermodynamic reference state

In thermodynamics, *ΔG* = *G*_bound_ − *G*_unbound_, where the unbound state includes the solvation energy of the free molecules, the entropy of the unbound system, and the standard-state concentration convention (1 M). None of these are represented in the SFM embedding. The intercept *b* absorbs all of this — it is the zero point of the energy scale, anchoring embedding-space scores to the experimental reference frame.

### S4.7 Temperature Recovery

The convergence equation predicts that the SFM’s learned temperature parameter *τ*_learned_ should correspond to the physical thermal energy *k*_*B*_*T* (or *RT* in molar units). This yields a specific, testable prediction.

**Experiment A (post-***τ*). Regress the post-temperature contrastive score *s* = ⟨**z**_agent_, **z**_target_⟩/*τ*_learned_ against *ΔG*. The slope *w*_*A*_ absorbs only the embedding-to-kcal/mol conversion.

**Experiment B (pre-***τ*). Regress the raw dot product *s*′ = ⟨**z**_agent_, **z**_target_⟩ (no temperature scaling) against *ΔG*. The slope *w*_*B*_ absorbs both the temperature and the unit conversion.

The Boltzmann identity predicts:

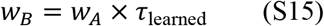

If this relationship holds empirically, the learned *τ* is functioning as the physical temperature parameter. This is temperature recovery: the model trained only on binary binding pairs (binder / non-binder) learns a continuous temperature parameter that corresponds to the physical *k*_*B*_*T*.

### S4.8 The Decoupling Principle

The affinity calibration framework decouples the SFM’s two data requirements:

#### The contrastive axis (cheap data, scaling)

Binary binding pairs — agent-target pairs known to bind, without measured affinities — train the contrastive embedding. These are available from co-crystal structures, display screening, FACS sorting, single-cell sequencing, and pseudo-labeling by structural models. The embedding geometry (and therefore the quality of the energy landscape) improves with more contrastive data. Affinity prediction quality improves correspondingly, even though the calibration data has not changed: a better energy landscape with the same unit conversion yields better absolute *ΔG* predictions.

#### The calibration axis (expensive data, saturating)

Measured *K*_*d*_ values — requiring purified proteins and SPR/ITC/BLI instrument time — provide the calibration targets. Because only two parameters (slope and intercept) are being fitted, the calibration regression is fully determined with very few measurements. Adding more *K*_*d*_ values beyond approximately 100–500 should yield diminishing returns — the limiting factor is the embedding geometry, not the unit conversion.

The **crossover point** — where adding more calibration data stops improving predictions but adding more contrastive data still helps — is the empirical signature of the decoupling. If observed, it establishes that the scaling axis for affinity prediction is the same as the scaling axis for retrieval: cheap binding pairs, not expensive *K*_*d*_ measurements.

### S4.9 Cross-Domain Calibration

The decoupling principle applies to every SFM domain where affinity data is scarce but binding data is abundant. The **leverage ratio** — contrastive pairs per calibration measurement — varies by domain:

| SFM | Contrastive data source | Calibration data source | Estimated leverage |
| --- | --- | --- | --- |
| CALM | SAbDab co-crystals + display | SAbDab $K_d$ annotations | 500:1 at scale |
| TCR-SFM | VDJdb + 10x multimer staining | Published TCR-pMHC $K_d$ | 200–500:1 |
| tSFM | JASPAR + ENCODE ChIP-seq | Published TF-DNA $K_d$ (PBM, HT-SELEX) | 100–1,000:1 |
| miR-SFM | miRTarBase + predicted targets | Measured duplex $\Delta G$ (NN model) | Given (exact) |
| eSFM | BRENDA + KEGG | BRENDA measured $K_m$ | 2–50:1 |
| dtSFM | PDBbind + BindingDB | PDBbind $K_d/K_i$ | 1.5–3:1 |
| mhcSFM | IEDB binary binding | IEDB quantitative $IC_{50}$ | 250:1 |
| crisprSFM | GUIDE-seq + CHANGE-seq | Measured gRNA-DNA $K_d$ | 1,000–10,000:1 |

Domains with the highest leverage (crisprSFM) benefit most from the decoupling: millions of cheap binding pairs, hundreds of expensive affinity measurements. Domains with low leverage (dtSFM, eSFM) have overlapping cheap and expensive data sources, and the decoupling advantage is smaller. For nucleic acid domains (miR-SFM, crisprSFM), the nearest-neighbor thermodynamic model provides exact *ΔG* values (SantaLucia, *PNAS*, 1998), making the “calibration” effectively free — the leverage ratio is infinite.

### S4.10 Verification Protocol

The following protocol tests the affinity calibration framework using existing SFM infrastructure:

**Step 1**. From the frozen SFM encoders, compute contrastive scores *s*_*ij*_ for all agent-target pairs with measured *K*_*d*_ values. Convert *K*_*d*_ to *ΔG* via Equation S12.

**Step 2**. Fit *ΔG* = *ws* + *b* (2 parameters) by ordinary least squares. Evaluate by Pearson *r*, Spearman *ρ*, RMSE, and MAE under 5-fold cross-validation with leakage-controlled splits.

**Step 3**. Fit *ΔG* = *as*^2^ + *ws* + *b* (3 parameters). Compare to the linear model by BIC. If the linear model is preferred (*Δ*BIC > 6), two-parameter sufficiency is confirmed.

**Step 4**. Repeat Steps 1–3 using the pre-*τ* raw dot product. Verify the temperature recovery prediction (Equation S15): *w*_*B*_ = *w*_*A*_ × *τ*_learned_.

**Step 5**. Repeat Steps 1–3 using a baseline encoder (e.g., ESM-2 without contrastive training). The gap between the SFM and the baseline quantifies the contribution of contrastive co-embedding to affinity prediction.

**Step 6**. Vary calibration set size (25, 50, 100, 250, 500 *K*_*d*_ values) at fixed contrastive training. Plot affinity *R*^2^ versus calibration set size. If the curve saturates by ∼100–250 points, the decoupling principle is confirmed.

### S4.11 Verification Checklist

□ Boltzmann factor chain: *K*_*d*_ = *k*_off_/*k*_on_ → *ΔG* = *RT*ln(*K*_*d*_) → *s*_*ij*_ = −*ΔG*_*ij*_/*RT* + *C*
□ Two-parameter sufficiency derived from convergence equation (slope = unit conversion, intercept = reference state)
□ Linear model preferred over quadratic by BIC (*Δ*BIC > 6)
□ Slope *w* is negative (higher similarity = tighter binding) and encodes kcal/mol per embedding unit
□ Intercept *b* absorbs thermodynamic reference state (solvation, entropy, standard-state concentration)
□ Temperature recovery: *w*_*B*_ = *w*_*A*_ × *τ*_learned_ holds empirically
□ Decoupling: affinity *R*^2^ saturates as function of calibration set size but continues improving with contrastive data
□ Leverage ratio estimated per domain
□ Baseline comparison (ESM-2) quantifies contrastive co-embedding contribution

## Supplementary Note S5

### Hybrid Recursive Learning: Cross-Modal Reinforcement Learning with Orthogonal Verification

#### Purpose

This note provides the mathematical formalization of the hybrid recursive learning framework introduced in §8 of the main text. We derive the actor-verifier decomposition from the convergence equation, formalize the orthogonal verification principle, specify the GRPO update rule for molecular design, derive the four collapse resistance mechanisms, and specify verifier architectures across all ten SFM domains. Every claim is explicit to enable verification.

### S5.1 Notation

| Symbol | Definition | Domain |
| --- | --- | --- |
| $\pi_\theta$ | Actor policy (SFM decoder with parameters $\theta$) | $\pi_\theta: \mathcal{C} \rightarrow \mathcal{X}$ |
| $\pi_{\text{ref}}$ | Frozen reference policy (pre-RL SFM decoder) | Same architecture as $\pi_\theta$ |
| $V(x, c)$ | Verifier score for candidate $x$ given target context $c$ | $V: \mathcal{X} \times \mathcal{C} \rightarrow \mathbb{R}$ |
| $\tau_{\text{RL}}$ | RL temperature (reward scaling) | $\tau_{\text{RL}} > 0$ |
| $\beta$ | KL penalty coefficient | $\beta > 0$ |
| $G$ | Group size (candidates per target) | $G \geq 2$ |
| $\hat{A}_i$ | Advantage of candidate $i$ within its group | $\hat{A}_i \in \mathbb{R}$ |
| $\epsilon_{\text{actor}}$ | Systematic error distribution of the actor | — |
| $\epsilon_{\text{verifier}}$ | Systematic error distribution of the verifier | — |

### S5.2 The Actor-Verifier Decomposition

The hybrid recursive learning framework decomposes molecular optimization into two components operating in orthogonal representation spaces.

#### Actor

The SFM decoder (§3.7 of the main text) proposes candidate molecular sequences *x* ∼ *π*_*θ*_(⋅ |*c*) conditioned on a target context *c* (the target embedding from the frozen dual encoder). The actor operates in sequence space: it generates discrete token sequences through either autoregressive decoding or discrete diffusion (Austin *et al., NeurIPS*, 2021; Sahoo *et al., arXiv*, 2024). The actor’s representation is the learned contrastive embedding — a vector space where the convergence equation governs selection probabilities.

#### Verifier

A structure-prediction or thermodynamic model evaluates each candidate *x* in a representation space orthogonal to the actor’s. For protein domains, the verifier is a co-folding model — AlphaFold3 (Abramson *et al., Nature*, 2024), Boltz-2 (Wohlwend *et al*., 2025), or an SE(3)-equivariant diffusion model — that generates structural ensembles and returns interface metrics. For nucleic acid domains, the verifier is a thermodynamic calculator — ViennaRNA (Lorenz *et al., Algorithms Mol. Biol*., 2011) or NUPACK (Zadeh *et al., J. Comput. Chem*., 2011) — that computes the exact Boltzmann energy *ΔG*.

The actor and verifier model the same physical quantity — binding free energy — through different mathematical machinery. The actor learns −*ΔG*/*k*_*B*_*T* through contrastive co-embedding and conditional sequence generation over token sequences. The verifier estimates *ΔG* through geometric interface energy computation (protein domains) or nearest-neighbor thermodynamic calculation (nucleic acid domains). The convergence equation governs both: the actor implements it as the training objective (InfoNCE, Supplementary Note S3); the verifier computes the physical quantity that the equation references.

### S5.3 The GRPO Update Rule for Molecular Design

The policy is updated using Group Relative Policy Optimization (GRPO; DeepSeek-AI, *arXiv*, 2025), adapted for molecular sequence generation.

#### Candidate generation

For each target context *c*, the actor generates a group of *G* candidate sequences: {*x*_1_, *x*_2_, …, *x*_*G*_} ∼ *π*_*θ*_(⋅ |*c*).

#### Verification

The verifier scores each candidate: *r*_*i*_ = *V*(*x*_*i*_, *c*) for *i* = 1, …, *G*. The score *r*_*i*_ is a scalar reward derived from the verifier’s output — for protein domains, a composite of interface metrics (ipTM, buried surface area, hydrogen bond count, clash score); for nucleic acid domains, the computed *ΔG*.

#### Advantage computation

Within each group, the advantage of candidate *i* is:

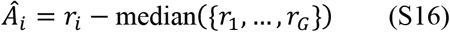

The median (rather than mean) provides robustness to outliers — a single candidate with an extreme verifier score does not dominate the group. Group-relative advantages eliminate the need for a learned value function (critic), removing a common source of instability in actor-critic methods (Schulman *et al., arXiv*, 2017).

#### Policy update

The GRPO objective is:

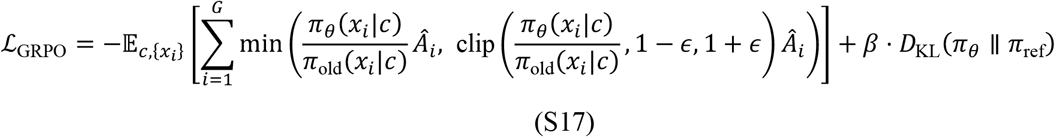

The first term is the clipped surrogate objective from PPO (Schulman *et al., arXiv*, 2017), applied with group-relative advantages. The second term is the KL divergence penalty to the frozen reference policy *π*_ref_, which preserves the actor’s base competencies (sequence fluency, diversity, physical plausibility). The clipping parameter *ε* (typically 0.1–0.2) prevents large policy updates, and the KL coefficient *β* is annealed during training — high initially to preserve the base model, reduced as the verifier signal is confirmed reliable.

#### No learned critic

Standard actor-critic methods (PPO, A2C) train a value network to estimate expected return, introducing its own approximation errors and instability. GRPO eliminates the critic entirely: advantages are computed from observed rewards within each group, not from a learned value estimate. DeepSeek-R1 demonstrated that this is sufficient for stable RL at scale when the verifier is reliable (DeepSeek-AI, *arXiv*, 2025).

### S5.4 The Orthogonal Verification Principle

#### Definition

Two models have orthogonal inductive biases if they share no architectural components, no training data, no loss functions, and no representational primitives.

#### Claim

The reliability of synthetic verification scales with the decorrelation of systematic errors between actor and verifier.

#### Formal statement

Let *ϵ*_actor_(*x*) be the systematic error of the actor’s implicit energy estimate for candidate *x*, and *ϵ*_verifier_(*x*) be the systematic error of the verifier’s energy estimate. In same-modality verification (actor and verifier share representational structure):

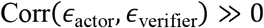

because shared representations propagate shared biases. In cross-modal verification (orthogonal inductive biases):

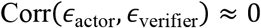

because the two representations have no structural overlap.

#### Consequence for reward hacking

Reward hacking occurs when the actor generates candidates that exploit systematic biases in the reward signal — candidates that score well according to the bias, not according to the physical truth. In same-modality verification, the actor can learn to exploit shared biases because high-reward regions of the actor’s representation correlate with high-reward regions of the verifier’s representation. In cross-modal verification, exploiting the verifier requires simultaneously exploiting two decorrelated bias distributions — a strictly harder optimization problem. The probability of successful reward hacking scales with Corr(*ϵ*_actor_, *ϵ*_verifier_): lower correlation means higher resistance.

#### Consequence for agreement reliability

When two models with correlated errors agree, the agreement may reflect shared bias rather than physical truth. When two models with uncorrelated errors agree, the agreement is evidence of physical truth — the probability that two independently biased systems produce the same incorrect answer is the product of their individual error probabilities. The orthogonal verification principle states that cross-modal agreement is more informative than same-modal agreement.

#### The SFM actor-verifier pair satisfies orthogonality

The sequence-space actor (autoregressive or discrete diffusion transformer; Vaswani *et al., NeurIPS*, 2017) and the structure-space verifier (SE(3)-equivariant diffusion model; Fuchs *et al., NeurIPS*, 2020) share no architectural components (transformer decoder vs. equivariant diffusion), no training data (binding pairs vs. PDB structures), no loss functions (InfoNCE vs. denoising score matching), and no representational primitives (token embeddings vs. 3D coordinates). Their error modes are decorrelated by construction.

### S5.5 Collapse Resistance: Four Independent Mechanisms

Policy collapse — convergence to a narrow, non-diverse distribution of high-reward outputs — is the primary failure mode of reinforcement learning with synthetic rewards (Amodei *et al., arXiv*, 2016). The hybrid recursive learning framework provides four independent collapse resistance mechanisms.

#### Mechanism 1: Error decorrelation (representational)

Cross-modal orthogonality (S5.4) raises the bar for reward hacking. To exploit the reward signal, the actor must generate candidates that simultaneously fool two fundamentally different mathematical frameworks — autoregressive or discrete diffusion token likelihood and SE(3)-equivariant geometric energy. This is strictly harder than fooling a single-modality verifier.

#### Mechanism 2: KL regularization (distributional)

The KL divergence penalty *β* ⋅ *D*_KL_(*π*_*θ*_ ∥ *π*_ref_) in Equation S17 constrains the policy to remain close to the reference distribution. This preserves the base competencies of the pre-RL actor: sequence fluency, germline proximity (for antibody domains), thermodynamic plausibility (for nucleic acid domains), and output diversity. The penalty is strongest at the beginning of training and is annealed as the reward signal is confirmed reliable. This is standard in RLHF (Ouyang *et al., NeurIPS*, 2022) and was shown to be sufficient for stability in verifier-only RL (DeepSeek-AI, *arXiv*, 2025).

#### Mechanism 3: Group-relative optimization (algorithmic)

GRPO computes advantages relative to the group median (Equation S16), eliminating the learned critic. A learned critic introduces its own approximation errors and can amplify reward hacking if its value estimates are biased. By computing advantages directly from observed rewards within each group, GRPO removes this source of instability. Within-group normalization also reduces variance: each update is relative to the current batch, not to a learned baseline that may drift.

#### Mechanism 4: Verifier abstention (signal quality)

When the verifier’s confidence falls below a threshold — measured by ensemble agreement for co-folding models, or by convergence diagnostics for thermodynamic calculators — it abstains: the candidate receives a reward of zero rather than a noisy estimate. This prevents the actor from learning on unreliable signals and bounds the noise in the reward surface. Calibrated abstention is essential: an uncalibrated verifier that always returns a score provides a reward surface the actor can exploit (Guo *et al., ICML*, 2017). The abstention rate is a diagnostic: a rising abstention rate indicates that the actor is moving into regions of sequence space where the verifier cannot reliably evaluate, which may be the first sign of distribution shift.

The four mechanisms operate at different levels — representational, distributional, algorithmic, and signal quality — providing defense in depth against collapse.

### S5.6 Domain-Specific Verifier Specifications

The verifier is the domain-dependent component of hybrid recursive learning. For each SFM domain, the appropriate verifier is determined by the available structural or thermodynamic models.

#### Protein-protein interaction domains (CALM, TCR-SFM, mhcSFM, rlSFM, rbpSFM)

Verifier class: co-folding diffusion models. Available models: AlphaFold-Multimer (Evans *et al., bioRxiv*, 2022), AlphaFold3 (Abramson *et al., Nature*, 2024), RoseTTAFold All-Atom (Krishna *et al., Science*, 2024), Boltz-1/Boltz-2 (Wohlwend *et al*., 2024, 2025), Chai-1 (Chai Discovery, 2024). Reward signal: composite of interface predicted TM score (ipTM), interface predicted aligned error (ipAE), buried surface area, hydrogen bond count, salt bridge count, and clash score. Abstention criterion: ensemble disagreement across multiple random seeds exceeds a threshold (e.g., ipTM standard deviation > 0.15 across 5 seeds). Orthogonality to actor: the co-folding model operates on 3D atomic coordinates via SE(3)-equivariant diffusion; the actor operates on token sequences via autoregressive or discrete diffusion decoding.

#### Nucleic acid recognition domains (tSFM, miR-SFM, crisprSFM)

Verifier class: thermodynamic free energy calculators. Available models: ViennaRNA (Lorenz *et al., Algorithms Mol. Biol*., 2011), NUPACK (Zadeh *et al., J. Comput. Chem*., 2011). Reward signal: computed duplex or secondary structure free energy *ΔG* using experimentally parameterized nearest-neighbor models (SantaLucia, *PNAS*, 1998). For crisprSFM, the reward also includes position-dependent mismatch penalties reflecting the seed-to-distal tolerance gradient (Hsu *et al., Nat. Biotechnol*., 2013). Abstention criterion: the computed *ΔG* is outside the parameterized range of the nearest-neighbor model (e.g., extreme GC content, non-canonical base pairs). Orthogonality to actor: the thermodynamic calculator operates on explicit energy parameters from calorimetric measurements; the actor operates on learned token embeddings. The verification signal in these domains is theoretically tighter than in protein domains because the verifier computes the exact Boltzmann energy, not a structural proxy.

#### Small molecule domains (eSFM, dtSFM)

Verifier class: molecular docking and binding free energy calculators. Available models: AutoDock Vina (Trott & Olson, *J. Comput. Chem*., 2010), GNINA (McNutt *et al., J. Cheminform*., 2021), DiffDock (Corso *et al., ICLR*, 2023). For higher accuracy at greater computational cost: free energy perturbation methods (FEP+; Wang *et al., J. Am. Chem. Soc*., 2015). Reward signal: docking score or computed *ΔG*_bind_. Abstention criterion: docking fails to converge, or the top-ranked pose has poor geometric complementarity (e.g., steric clashes exceeding a threshold). Orthogonality to actor: the docking model operates on 3D molecular coordinates and force-field-derived energies; the actor operates on SMILES or molecular graph token sequences.

### S5.7 Relationship to Standard RLHF

The standard RLHF framework (Ouyang *et al., NeurIPS*, 2022) trains a reward model from human preference data, then uses the reward model to optimize the policy via PPO. Hybrid recursive learning is a special case of the broader reward-model-free RL paradigm, where the reward model is replaced by a physics-based verifier. The key differences are:

| Component | Standard RLHF | Hybrid recursive learning |
| --- | --- | --- |
| Reward source | Human preferences | Physics-based verifier |
| Reward model | Learned from preference data | Not needed (verifier is the reward) |
| Critic / value network | Learned (PPO) | Not needed (GRPO) |
| Actor-reward correlation | High (shared training data, same modality) | Low (orthogonal representations) |
| Abstention | Not standard | Built-in (ensemble disagreement) |
| Domain specificity | General (language) | Domain-specific (per SFM) |

The hybrid recursive learning framework inherits the GRPO innovation from DeepSeek-R1 — eliminating both the learned reward model and the learned critic — and adds the orthogonal verification principle as a theoretical basis for reward signal reliability.

### S5.8 Scope and Limitations

#### What the theory establishes

The orthogonal verification principle provides a theoretical basis for expecting cross-modal actor-verifier pairs to be more robust against reward hacking than same-modal pairs. The GRPO update rule with KL regularization and verifier abstention provides four independent collapse resistance mechanisms. The framework generalizes across all ten SFM domains.

#### What the theory does not establish

The framework does not guarantee convergence to optimal policies, nor does it specify the rate of improvement. The orthogonal verification principle is a qualitative argument about error decorrelation, not a quantitative bound on reward hacking probability. The verifier’s accuracy is assumed, not proven — if the structural verifier is systematically wrong (e.g., co-folding models that produce physically unrealistic structures for designed sequences, as discussed in §9.5 of the main text), the RL loop will optimize toward verifier artifacts rather than physical truth. Periodic wet-lab validation is necessary to confirm that computational gains translate to experimental binding.

#### Implementation dependencies

The framework requires: (i) a trained SFM decoder capable of conditional sequence generation, (ii) a calibrated verifier with reliable abstention, (iii) sufficient compute for generating and verifying groups of candidates per target. The compute requirement scales as *G* × *T*_verify_ per target per iteration, where *T*_verify_ is the verifier’s per-candidate evaluation time. For co-folding verifiers, this is minutes per candidate; for thermodynamic calculators, milliseconds.

### S5.9 Verification Checklist

□ Actor-verifier decomposition: actor in sequence space, verifier in structure/energy space
□ GRPO update rule specified (Equation S17): clipped surrogate + KL penalty, no learned critic
□ Advantages computed relative to group median (Equation S16), robust to outliers
□ Orthogonal verification principle: decorrelated errors → higher reward signal reliability
□ Four collapse resistance mechanisms at four levels: representational, distributional, algorithmic, signal quality
□ Verifier specified per domain: co-folding for proteins, thermodynamic calculators for nucleic acids, docking for small molecules
□ Abstention criterion defined per verifier class
□ Relationship to standard RLHF clarified: hybrid RL is reward-model-free, critic-free
□ Limitations acknowledged: no convergence guarantee, verifier accuracy assumed, wet-lab validation required

## Notes

### Competing Interest Statement

S.T.R. is a co-founder and scientific advisor of Engimmune Therapeutics, Encelta, Fy Cappa Biologics and BIIE.AI.

### Funding

- Foundation Immune Engineering for Global Child and Adolescent Health (FIE).: Vine

